# Spatio-temporal plasticity of root exudation in three temperate tree species: effects of season, site and soil characteristics

**DOI:** 10.1101/2025.09.21.675509

**Authors:** Melissa Wannenmacher, Simon Haberstroh, Jürgen Kreuzwieser, Jörg Niederberger, Jörg Prietzel, Friederike Lang, Christiane Werner

## Abstract

Root exudation provides a constant carbon input to the rhizosphere and is therefore a very important factor in shaping this hotspot of biological activity. Nonetheless, root exudation data and its spatio-temporal plasticity is scarce. This study provides insights into compound-specific root exudation in three temperate tree species in two seasons (late spring and late summer) and two soil compartments (forest floor and the top mineral soil), including the effect of soil chemistry. At four sites with differing mean annual temperature and soil phosphorus level, root exudates were sampled using an *in-situ* cuvette-based system and analysed by gas chromatography-mass spectrometry. We found seasonally and spatially varying site- and species-specific exudation patterns. While the seasonal pattern was similar among species and sites, with higher exudation rates in late spring, soil compartment-specific exudation depended on species and site. *Acer pseudoplatanus* tended to exude more into the mineral soil at warmer sites, while *Picea abies* exuded more in the mineral soil at all sites. Exudation by *Fagus sylvatica* was independent from the soil compartment. Significant correlation between exchangeable soil cations and specific compounds exuded by *F. sylvatica* and *P. abies* were found. Exudation of specific compounds in *F. sylvatica* increased with the concentration of exchangeable Mg, Al and Fe, whereas exudation rates in *P. abies* decreased with most base cations’ concentration, while sugar exudation increased with the exchangeable non-base cations Al and Fe. These results demonstrate that root exudation is dynamically adjusted to the species-specific nutritional needs governed by site, season and soil characteristics.

## Introduction

Root exudates play a major role in carbon and nutrient cycling of forests and transform the rhizosphere into a hub of biological activity. Various definitions of the term root exudation can be found, which comprise different selections of compounds. Most commonly, it includes organic compounds released by the living plant root, which distinguishes it from rhizodeposition more commonly seen as compounds set free by dead plant roots (Massalha *et al*., 2017). In this study, we use the term root exudates for soluble chemical compounds, which are released by living plant roots. Numerous functions of root exudates have been identified including defence against pathogens, attraction of beneficial microbes and plant-plant-communication (Dannenmann *et al*., 2009; Huang *et al*., 2014; Rohrbacher & St-Arnaud, 2016). Through the so-called priming effect, plants make organically bound nutrients bioavailable by providing easily degradable carbon (C) compounds to microorganisms as energy source to decompose more recalcitrant components in the soil (Rohrbacher & St-Arnaud, 2016; Tückmantel *et al*., 2017). Ruf *et al*. (2006) state that root exudates are, next to litter and soil organic matter (SOM), one of the three primary C sources in the soil food web, which even reach the third trophic level, such as predatory mites. Moreover, rhizodeposition, even in small amounts, is responsible for up to one third of C- and nitrogen (N)-mineralisation in temperate forest soils (Finzi *et al*., 2015).

Next to the priming effect, plants are able to exude compounds, which directly increase the mobility and bioavailability of nutrients (Rohrbacher & St-Arnaud, 2016), demonstrating that nutrient uptake and root exudation are closely interrelated. However, there is no clear pattern on how nutrient availability influences root exudation and how root exudation rate and quality mediate nutrient uptake (Williams *et al*., 2022). While generally decreased exudation under phosphorus (P) limiting conditions has been reasoned by decreased meristematic activity on the one side (Canarini *et al*., 2019), other studies found increased exudation of organic acids when P was limiting (Jones *et al*., 2009; Vives-Peris *et al*., 2020). These organic acids have been identified to help mobilise P before (Jones *et al*., 2004). This effect was even greater under acidic conditions (Jones *et al*., 2004). The same effect was found for iron (Fe) (Jones *et al*., 2004). Supporting this, increased exudation of organic acids under Fe limiting conditions were found by Vives-Peris *et al*. (2020), coupled with an increase of exudation of sugars (Vives-Peris *et al*., 2020) and phenolics (Vranova *et al*., 2013). A decrease and increase of 2-benzopyranons exudation under P and Fe limiting conditions, respectively, has also been observed (Pantigoso *et al*., 2021). Contrasting those findings, a recent study on beech suggests that there is no effect of P availability on root exudation (Leuschner *et al*., 2022). However, this latter study examined the total exuded C not looking at organic acids specifically. Hence, the increased organic acid exudation under P limiting conditions might be compensated by a reduction of exudation in other compound groups. Exchangeable aluminium (Al^3+^) abundance also seems to increase the exudation of organic acids (Jones *et al*., 2009; Canarini *et al*., 2019). Hereby, malate, pyruvate and citrate have been identified to help withstand Al toxicity (Pantigoso *et al*., 2021). The effect of other nutrients on root exudation is less studied. Nonetheless, it has been suggested that deficits of boron (B), zinc (Zn) and manganese (Mn) lead to increased exudation due to a loss of membrane functioning (Vranova *et al*., 2013). Further, potassium (K) starvation has been found to inhibit the exudation of sugars (Vives-Peris *et al*., 2020). Next to the nutrient composition, more general site conditions also impose an effect on root exudation. In beech, a tripling of root exudation C rate was observed, when the acidity doubled (Meier *et al*., 2020). Another study on *Lactuca sativa* did not find such a stringent relationship with pH, but identified the soil type as an important factor influencing root exudation (Neumann *et al*., 2014). Salt stress for example, also led to higher exudation of specific compounds in Arabidopsis and walnut (Vives-Peris *et al*., 2020; Pantigoso *et al*., 2021). Most of the above-mentioned studies were conducted on crops and grasses. Despite their importance, only little is known about root exudation rates, composition and strategies in different temperate tree species across seasons and soil compartments (Möller *et al*., 2024). The fewer studies on trees might be due to the more challenging sampling procedure, especially when working in forests. Compared to grasses, trees use a relatively smaller share of photoassimilates for root exudates (Vives-Peris *et al*., 2020), which seems to vary with species and site condition ranging from 0.6% (Brunn *et al*., 2022) (in beech) to 5% (Phillips *et al*., 2011) (in loblolly pine) according to previous studies. Nonetheless, root exudation of trees is an important factor for the forest environment.

Root exudates, for instance, play a major role in shaping the composition and thickness of the forest floor (Prescott & Grayston, 2013; Finzi *et al*., 2015). In the present work, the forest floor is defined as the upper-most layer of the forest soil, which mainly consists of litter in various stages of decomposition or transformation with a minimum of 15% organic C in mass (Wachendorf *et al*., 2023) and therefore represents a stock of organically bound nutrients. The forest floor has been identified as an important nutrient source for forest trees, especially at nutrient-poor sites (Lang *et al*., 2017). In line with this, indications that the presence of organic N, as present in the forest floor, increases exudation have been found (Rohrbacher & St-Arnaud, 2016; Tückmantel *et al*., 2017; Vives-Peris *et al*., 2020). Tückmantel *et al*. (2017) found that ectomycorrhiza (EM) associated tree species exuded more C compared to arbuscular mycorrhiza (AM) associated species. EM-tree associations are able to produce extracellular enzymes and follow an organic nutrient economy, while AM-tree associations often lack this feature and follow an inorganic nutrient economy (Phillips *et al*., 2013; DeForest & Snell, 2020). The inorganic nutrient economy is characterised by a rapid turn-over of the AM plant litter with a comparatively high quality. In contrast, EM plant litter has a lower quality, which leads to a slower turn-over and therefore a larger nutrient storage pool in the organic layer. Hence, in theory, EM-associated tree species should show a higher specialisation on nutrient uptake from organic soil material relying on a tighter nutrient recycling in the forest compared to AM-associated tree species, which rather acquire nutrients from mineral sources. Since different tree species follow different nutrient uptake strategies, it is likely that they also follow different root exudation strategies. The source of nutrients in forest floors is mainly organic matter while in the mineral soil, inorganic sources play a bigger role, which could be mirrored in the quality and quantity of root exudates in the different tree species.

A seasonally varying nutrient demand could also be reflected in seasonal exudation patterns. Strong growth in spring increases the plant’s nutrient demand, while during the period before dormancy in late summer and autumn a smaller amount is needed (Carrara *et al*., 2004). At the same time, photosynthetic activity is higher in spring allowing for C investment in root exudates. However, Tückmantel *et al*. (2017) did not find variation in exudation across seasons (within one growing season) in beech. Also, the variation of exudation within days or weeks seems to be low (Phillips *et al*., 2008). On the other side, there are studies that found seasonal exudation patterns (Jakoby *et al*., 2020; Chen *et al*., 2023). These contradictory findings reflect the lack of knowledge in this field.

The main objectives of this study were to investigate (i) seasonal exudation patterns of the two EM-associated tree species European beech (*Fagus sylvatica*) and Norway spruce (*Picea abies*), and the AM-associated tree species Sycamore maple (*Acer pseudoplatanus*) (ii) in the forest floor and the upper 5 cm of the mineral soil at four temperate forest sites with differing abiotic conditions. Finally, we aimed at (iii) investigating the influence of soil chemistry on exudation patterns of the three species.

We hypothesise that trees can adjust their exudation (quantity and composition) on a temporal and a small spatial scale, such as between soil compartments, to their species-specific nutritional needs in dependence on the distribution of nutrient availability in the soil. We hereby expect the EM associated species (beech and spruce) to show a generally higher exudation, which is more dominant in the forest floor, compared to a generally lower and mineral soil focused exudation in the AM associated species (maple).

## Material and Methods

### Study sites

Samples were taken at four European beech (*Fagus sylvatica*)-dominated temperate forest sites. Waldkirch (WAL) and Kandel (KAN) are located in the Black Forest in southwest Germany. The site Kaiserstuhl (KAI) is situated in the same region, but in the upper Rhine valley. Bad Brückenau (BBR) lies in the Rhön mountains in Central Germany. Basic information on the sites can be found in Table 1. The sites mainly differ in mean annual temperature (MAT) and phosphorus (P) level in the soil (Tables 1 & 2). The two sites at lower altitudes WAL and KAI showed a higher MAT in the reference period 1991 – 2020 than the sites KAN and BBR at higher altitudes. Being located on basalt rock, KAI and BBR can be classified as P-rich, while the other two sites located on granite rock are P-poor (Table 2). The sites with higher MAT, KAI and WAL, are less acidic than the cooler sites. The soil type was classified according WRB 4^th^ edition (IUSS Working Group WRB, 2022), the humus form according the current German soil classification system (KA6) (Wachendorf *et al*., 2023). Mean annual temperature (MAT) and mean annual precipitation (MAP) were deducted from 1×1 km grid data HYDRAS-DE-TAS (for MAT) and HYDRAS-DE-PRE (for MAP) provided by the DWD Climate Data Center (CDC). According to the DWD, data are based on the period from 1991 to 2020. Standardised precipitation evaporation index (SPEI) values (Vicente-Serrano *et al*., 2010) for the sites were looked up at the SPEI database (https://spei.csic.es/spei_database/) (Beguería *et al*., 2010; Vicente-Serrano *et al*., 2010). The values calculated on the base of the six previous months were chosen here.

**Table 1:** Site information with coordinates, altitude, mean annual temperature (MAT), mean annual precipitation (MAP) from the years 1991 - 2020 (DWD), parent material, soil type and humus form.

| site | coordinates | altitude<br>(m.a.s.l.) | parent<br>material | soil type | humus form | MAT<br>[°C] | MAP<br>[mm] |
| --- | --- | --- | --- | --- | --- | --- | --- |
| **WAL** | 48.1° N,<br>7.9° E | 415 | Granite | Dystric Cambisol | Typical F-mull | 10.3 | 1107 |
| **KAI** | 48.1° N,<br>7.7° E | 396 | Basalt | Skeletal<br>Umbrisol | Typical F-mull<br>- typical moder | 9.3 | 984 |
| **KAN** | 48.1° N,<br>8.0° E | 1163 | Granite | Dystric Skeletal<br>Rhodic Cambisol | Typical moder | 6.3 | 1769 |
| **BBR** | 50.4° N,<br>9.9° E | 809 | Basalt | Eutric Cambisol | Typical F-mull<br>- typical moder | 6.8 | 1110 |

**Table 2:** Soil element concentration, exchangeable cations, base saturation and pH (H_2_O), at the four study sites Waldkirch (WAL), Kaiserstuhl (KAI), Kandel (KAN) and Bad Brückenau (BBR), in the forest floor and mineral soil.

|  | WAL |  | KAI |  | KAN |  | BBR |  |
| --- | --- | --- | --- | --- | --- | --- | --- | --- |
|  | forest floor | mineral soil | forest floor | mineral soil | forest floor | mineral soil | forest floor | mineral soil |
| Al [mg/g] | $20.1 \pm 4.0$ | $27.3 \pm 1.6$ | $13.1 \pm 2.5$ | $21.0 \pm 2.3$ | $16.2 \pm 0.7$ | $42.2 \pm 2.4$ | $14.3 \pm 4.8$ | $33.2 \pm 2.2$ |
| Ca [mg/g] | $6.1 \pm 0.6$ | $1.3 \pm 0.1$ | $18.2 \pm 1.4$ | $11.9 \pm 5.8$ | $2.2 \pm 0.1$ | $0.6 \pm 0.0$ | $9.0 \pm 1.2$ | $3.5 \pm 0.5$ |
| Fe [mg/g] | $9.0 \pm 1.7$ | $26.4 \pm 1.0$ | $14.6 \pm 2.0$ | $35.5 \pm 1.8$ | $10.8 \pm 1.2$ | $39.4 \pm 2.0$ | $17.4 \pm 5.4$ | $49.3 \pm 1.9$ |
| K [mg/g] | $17.4 \pm 2.3$ | $2.6 \pm 0.4$ | $8.3 \pm 2.1$ | $3.4 \pm 0.6$ | $5.2 \pm 1.0$ | $8.0 \pm 0.6$ | $4.5 \pm 1.4$ | $2.4 \pm 0.1$ |
| Mg [mg/g] | $1.1 \pm 0.0$ | $1.7 \pm 0.3$ | $4.6 \pm 0.8$ | $5.3 \pm 1.9$ | $1.9 \pm 0.1$ | $4.4 \pm 0.3$ | $3.3 \pm 0.5$ | $5.9 \pm 0.7$ |
| Mn [mg/g] | $4.2 \pm 0.5$ | $3.0 \pm 0.2$ | $1.3 \pm 0.1$ | $0.7 \pm 0.2$ | $0.2 \pm 0.0$ | $0.2 \pm 0.0$ | $1.1 \pm 0.1$ | $1.2 \pm 0.1$ |
| Na [mg/g] | $1.5 \pm 0.5$ | $0.3 \pm 0.0$ | $2.4 \pm 0.8$ | $0.3 \pm 0.0$ | $0.8 \pm 0.4$ | $0.2 \pm 0.0$ | $1.0 \pm 0.5$ | $0.3 \pm 0.0$ |
| P [mg/g] | $0.7 \pm 0.0$ | $0.5 \pm 0.0$ | $1.2 \pm 0.1$ | $1.4 \pm 0.1$ | $0.9 \pm 0.1$ | $0.9 \pm 0.0$ | $1.3 \pm 0.2$ | $2.1 \pm 0.1$ |
| Al<sup>3+</sup> [μmol/g] | $0.8 \pm 0.2$ | $8.7 \pm 2.0$ | $0.6 \pm 0.3$ | $8.9 \pm 6.6$ | $21.0 \pm 11.5$ | $77.2 \pm 12.7$ | $12.7 \pm 6.1$ | $49.6 \pm 3.7$ |
| **Ca<sup>2+</sup>**<br>**[μmol/g]** | 225.4±27.3 | 28.3±5.4 | 304.6±22.6 | 100.1±44.6 | 207.3±57.9 | 16.7±8.1 | 167.5±28.3 | 48.2±7.5 |
| **Fe<sup>2+</sup>**<br>**[μmol/g]** | 0.1 ± 0.0 | 0.1 ± 0.0 | 0.1 ± 0.0 | 0.8 ± 0.7 | 2.3 ± 1.1 | 6.5 ± 1.5 | 0.9 ± 0.4 | 2.3 ± 0.4 |
| **K<sup>+</sup>**<br>**[μmol/g]** | 55.8 ± 1.8 | 4.4 ± 0.6 | 39.8 ± 5.8 | 4.7 ± 0.8 | 21.3 ± 3.0 | 4.4 ± 0.5 | 19.7 ± 2.9 | 6.9 ± 1.1 |
| **Mg<sup>2+</sup>**<br>**[μmol/g]** | 51.3 ± 8.9 | 5.4 ± 0.9 | 101.7 ± 4.2 | 21.7 ± 3.4 | 69.9 ± 11.9 | 10.6 ± 1.9 | 88.4 ± 16.1 | 25.3 ± 4.5 |
| **Mn<sup>2+</sup>**<br>**[μmol/g]** | 52.3 ± 8.8 | 16.9 ± 2.2 | 14.9 ± 2.7 | 2.4 ± 0.6 | 15.1 ± 4.6 | 1.3 ± 0.7 | 15.9 ± 1.7 | 6.7 ± 0.4 |
| **Na<sup>+</sup>**<br>**[μmol/g]** | 2.5 ± 0.6 | 0.3 ± 0.1 | 2.0 ± 0.4 | 0.9 ± 0.2 | 1.3 ± 0.2 | 0.7 ± 0.1 | 2.2 ± 0.6 | 1.2 ± 0.3 |
| **H<sup>+</sup>**<br>**[μmol/g]** | 0.3 ± 0.1 | 4.7 ± 0.0 | 0.8 ± 0.3 | 5.4 ± 3.5 | 32.6 ± 9.0 | 20.7 ± 3.9 | 9.7 ± 1.2 | 11.5 ± 1.1 |
| **CEC**<br>**[μmol/g]** | 389.8±23.2 | 67.9±5.7 | 465.2±30.4 | 142.6±43.2 | 370.9±56.9 | 138.1±8.2 | 317.1±42.2 | 151.7±12.5 |
| **BS** | 85.7 ± 3.6 | 55.3 ± 6.3 | 96.2 ± 0.8 | 83.2 ± 11.2 | 74.0 ± 10.1 | 25.8 ± 9.3 | 83.8 ± 4.9 | 51.6 ± 4.8 |
| **pH**<br>**(H<sub>2</sub>O)** | 5.9 ± 0.1 | 5.1 ± 0.0 | 6.3 ± 0.2 | 5.5 ± 0.5 | 4.4 ± 0.2 | 4.0 ± 0.1 | 5.1 ± 0.2 | 4.5 ± 0.0 |

Soil and forest floor pH (H_2_O) of air-dried samples (40°C) were determined using double deionized water at a ratio of 1:5 (soil:solution). Samples were analysed using Metrohm CH/789 Robotic Sample Processor XL, measuring chain: Micro El.Cone 16 WOC.

Exchangeable cation concentrations were determined using NH_4_Cl at a concentration of 0.5 mol l^-1^. The concentrations of extracted cations (Ca, Mg, K, Na, Mn, Al, Fe) were determined by inductively coupled plasma optical emission spectroscopy (IPC-OES 5800, Agilent, Santa Clara, USA). The proton concentration (used for CEC calculation) was determined by titration using 0.05 mol NaOH and with Metrohm CH, Titrando 905/789 Robotic Sample Processor XL, measuring chain: Micro El.Cone 16 WOC.

For the total soil element content analysis, the dried and milled soil and forest floor samples underwent HNO_3_ microwave pressure digestion at 170°C, followed by inductively coupled plasma atomic emission spectroscopy (IPC-OES 5800, Agilent, Santa Clara, USA).

A summary of soil element concentrations, pH, cation exchange capacity (CEC) and the base saturation (BS) in the soil compartments relevant for exudation sampling can be found in Table 2. The given values were averaged over three to five sampling points per site and soil compartment. The forest floor thickness was measured and averaged over three spots around each sampling tree with a distance of 1.5 m from the stem. Small holes were dug to obtain a flat vertical surface, where the forest floor – mineral soil interface was clearly visible. The forest floor thickness ranged from a minimum of 0.2 cm at WAL (late spring: 2.6 ± 2.0 cm (mean ± standard deviation), late summer: 2.1 ± 1.6 cm) over KAI (late spring: 2.1 ± 1.0 cm, late summer: 2.0 ± 1.8 cm) and BBR (late spring: 6.1 ± 1.7 cm, late summer: 4.2 ± 2.0 cm) to a maximum of 10.0 cm at KAN (late spring: 6.3 ± 1.7 cm, late summer: 3.9 ± 2.0 cm).

### Sampling design

Sampling took place on five trees per species per site in two campaigns in late spring and late summer in 2023. No spruce trees were present at KAI. The diameters at breast height (DBH) of the sampling trees ranged from 3.2 cm to 40.4 cm in maple, from 25.8 cm to 47.8 cm in beech and from 31.8 cm to 63.3 cm in spruce (Table 3). The smaller DBH in maple were due to the absence of thicker individuals of maple at the selected sites. The sampling campaigns lasted from 16^th^ of May to 6^th^ of June (KAI: 16/05 – 20/05, WAL 22/05 – 26/05, KAN: 28/05 – 01/06, BBR: 03/06 – 06/06) and from 29^th^ of August to 19^th^ of September (KAN: 29/08 – 02/09, WAL: 04/09 – 07/09, BBR: 10/09 – 13/09, KAI: 16/09 – 19/09). In all campaigns, root exudates from the same trees were sampled to avoid sampling artefacts through specific exudation patterns of individual trees. In each campaign, two roots per tree were investigated, one in the forest floor and one in the mineral soil. Additionally, between 7 and 10 control cuvettes per site without roots were exposed to the same sampling procedure, as described below.

**Table 3:** Mean DBH ± standard deviation at the four sites for the tree species *Acer pseudoplatanus* (maple), *Fagus sylvatica* (beech) and *Picea abies* (spruce).

|  | maple | beech | spruce |
| --- | --- | --- | --- |
| **WAL** | 13.9 ± 3.6 cm | 43.2 ± 4.0 cm | 44.8 ± 10.0 cm |
| **KAI** | 7.6 ± 5.3 cm | 36.5 ± 7.5 cm | - |
| **KAN** | 8.9 ± 3.7 cm | 33.5 ± 3.0 cm | 48.3 ± 6.7 cm |
| **BBR** | 20.7 ± 13.5 cm | 39.3 ± 7.1 cm | 38.5 ± 5.7 cm |

### Root exudate collection

An *in-situ* cuvette-based approach based on Phillips *et al*. (2008) was used for the exudate collection. Two roots per tree, one in the forest floor and one in the mineral soil were excavated and traced back to the tree to correctly identify the tree individuum and species. Soil particles were carefully removed with water and tweezers while the roots were still attached to the tree. Tap water was used for cleaning instead of distilled water to avoid stressing the root through a high osmotic potential within and outside the plant cells. Aluminium foil was wrapped around the cleaned root to let it recover from the excavation and cleaning process for 24 hours. The aluminium foil was covered with litter to avoid animals being attracted by it. After this recovery time, the root was briefly rinsed with tap water and then placed in a cuvette consisting of a 30 ml plastic syringe with Luer-Lock fitting (B.Braun, Melsungen, Germany) with a 3-way-valve (Teqler, Wecker, Luxembourg) attached to it. The syringe plunger was removed and the outlet was covered with glass wool. 3 mm glass beads and 10 ml of a C-free dilute nutrient solution (0.5 mM NH_4_NO_3_, 0.1 mM KH_2_PO_4_, 0.2 mM K_2_SO_4_, 0.15 mM MgSO_4_*7H_2_O, 0.4 mM CaCl_2_*2H_2_O) were added to imitate the soil environment. Afterwards, the cuvette was sealed with parafilm to avoid contamination. 24 hours later the nutrient solution was removed using a syringe. Two flushes with a C- and N-free dilute nutrient solution (0.1 mM KH_2_PO_4_, 0.2 mM K_2_SO_4_, 0.15 mM MgSO_4_*7H_2_O, 0.4 mM CaCl_2_*2H_2_O) were performed before adding 5 ml of the same C- and N-free solution as exudation capturing solution. After 24 hours, the capturing solution was removed and two flushes of 10 ml were performed to capture the exudates attached to the glass beads and the root. The retrieved solution was immediately filtered through a 0.2 µm sterile syringe filter. In the field, exudation solutions were kept in a cooling box and covered by frozen ice pads before being frozen to -80°C at the lab until further analysis. The control cuvettes were handled in the exact same way as the cuvettes containing roots. The roots in the cuvettes were cut, cooled at ∼8°C until scanning. The root surface area was determined using the software WinRHIZO 2021a 32-Bit (Regent Instruments Inc., Quebec, Canada) with a scanner (Perfection V850 Pro, Seiko Epson Corporation, Tokyo, Japan).

### Root exudate analysis with GC-MS

The frozen root exudate samples were lyophilised for ∼90 h and the dried exudates derivatised. Derivatisation was conducted as specified in Maurer *et al*. (2021). In brief, the lyophilised exudates were solved in 100 µl of a 20 mg ml^-1^ solution of methoxyamine hydrochloride in anhydrous pyridine. After centrifugation, 20 µl of the prepared solution were incubated on the thermoshaker at 30°C and 1400 rpm for 90 min. Then, 35 µl of N-methyl-N-(trimethylsilyl)-trifluoroacetamide (MSTFA) were added and samples were shaken at 37°C and 1400 rpm for 30 min. The derivatives were analysed by GC-MS (GC 7820 gas chromatograph coupled to a 5975C MSD, Agilent, Santa Clara, USA).

Parameter settings were the same as specified in Kreuzwieser *et al*. (2009) with a split of 2:1. For peak detection, identification, and peak area determination the MASSHUNTER Quantitative Analysis software (Agilent Technologies, Santa Clara, USA) with the Golm Metabolome Database (2011) was used. Only compounds with a match factor larger than 60 were included in the analysis. The identified compounds were classified into 11 groups based on their chemical properties: amino acids, organic acids, fatty acids (containing more than 7 carbon atoms), inorganic acids, alcohols, sugars, N-containing and N-free aromatic compounds, other N-containing compounds, other hydrocarbons and others, which comprises non-identified compounds.

### Quantification of exudation rates

For all analysed compounds, the peak area means of the controls from one sampling period and site were subtracted from the peak areas of the respective samples to account for possible contamination. Exudation rates were calculated based on the amount of the exuded compound per the root surface area and per exposure time of roots in the cuvette.

The following compounds were quantified using authentic standards: alanine, aspartic acid, boric acid, citric acid, cysteine, fructose, galactose, glutamic acid, glycerol, lactic acid, leucine, malic acid, N,N-dimethyl-glycine, phenylalanine, propane-1,3-diol, pyruvic acid, sucrose, triethanolamine and uric acid. The peak areas of those compounds covered > 75% of the total peak area in the samples. For the compounds not covered by the mentioned standards, a mean quantification factor was calculated from the standards within the same group. A list of the identified compounds, their classification and the standard used for quantification can be found in Supplementary Table S1.

Root exudation has been reported in different units, e.g. µmol C g_dry root_^-1^ h^-1^ when focussing on plant physiological processes and g C (10 cm)^-1^ m^-2^ a^-1^ when focussing on ecosystem carbon fluxes (Tückmantel *et al*., 2017). In this study, we chose the root surface area as a reference (ng cm^-2^ h^-1^), since it is a better predictor for root exudation (Jakoby *et al*., 2020) and represents the direct exchange layer between the root and the rhizosphere, where root exudates and nutrients are transferred. Furthermore, it allows for valid comparisons between species.

### Statistical analysis

All data were analysed using R (version 4.3.2, R Core Team 2024) in RStudio (version 2024.12.0). To determine significant differences between seasons and soil compartments, the Wilcoxon test was used, since normality of the residuals of the exudation data set was not given. The strength of correlations between compound groups in the exudation and the concentration of soil elements were determined with the Spearman’s rank correlation coefficient. To test for significances in those correlations, the function cor.test with the option “method = ‘spearman’” was used. Regressions between compound groups in the exudation and the concentration of soil elements were visualised using a generalised linear smoothing.

## Results

In total, 72 compounds detected by GC-MS were included in the analysis. The most abundant compounds were N,N-dimethylglycine, triethanolamine, boric acid, and octadecadienoic acid, which covered ∼60% of exudation mass included in the analysis. The most important compound groups were amino acids, other N-containing compounds and inorganic acids. Sugars and N-containing aromatic compounds were the least abundant compound groups; however, those groups showed a high responsiveness to environmental conditions. A full list of identified compounds, including the compound class can be found in Supplementary Table S1.

Amongst the tree species, maple and beech showed the highest exudation rates with 131 ± 30 ng cm^-2^ h^-1^ and 103 ± 23 ng cm^-2^ h^-1^ (mean across all samples and both seasons ± 1 standard error), respectively, while spruce exuded significantly less than maple (p ≤ 0.01) with 61 ± 11 ng cm^-2^ h^-1^ (Figure 1). Amino acids made up the highest share of exudation with 48%, 37% and 34% for maple, beech and spruce, respectively. The share of all N-containing compounds in the exudation of maple and beech made up more than one half with 60% and 54%, respectively, while for spruce they amounted for 47% of exudation. In contrast, spruce showed a higher share of non-N-hydrocarbons, such as aromatics, fatty acids and sugars compared to the other species (Figure 1).

**Figure 1:**
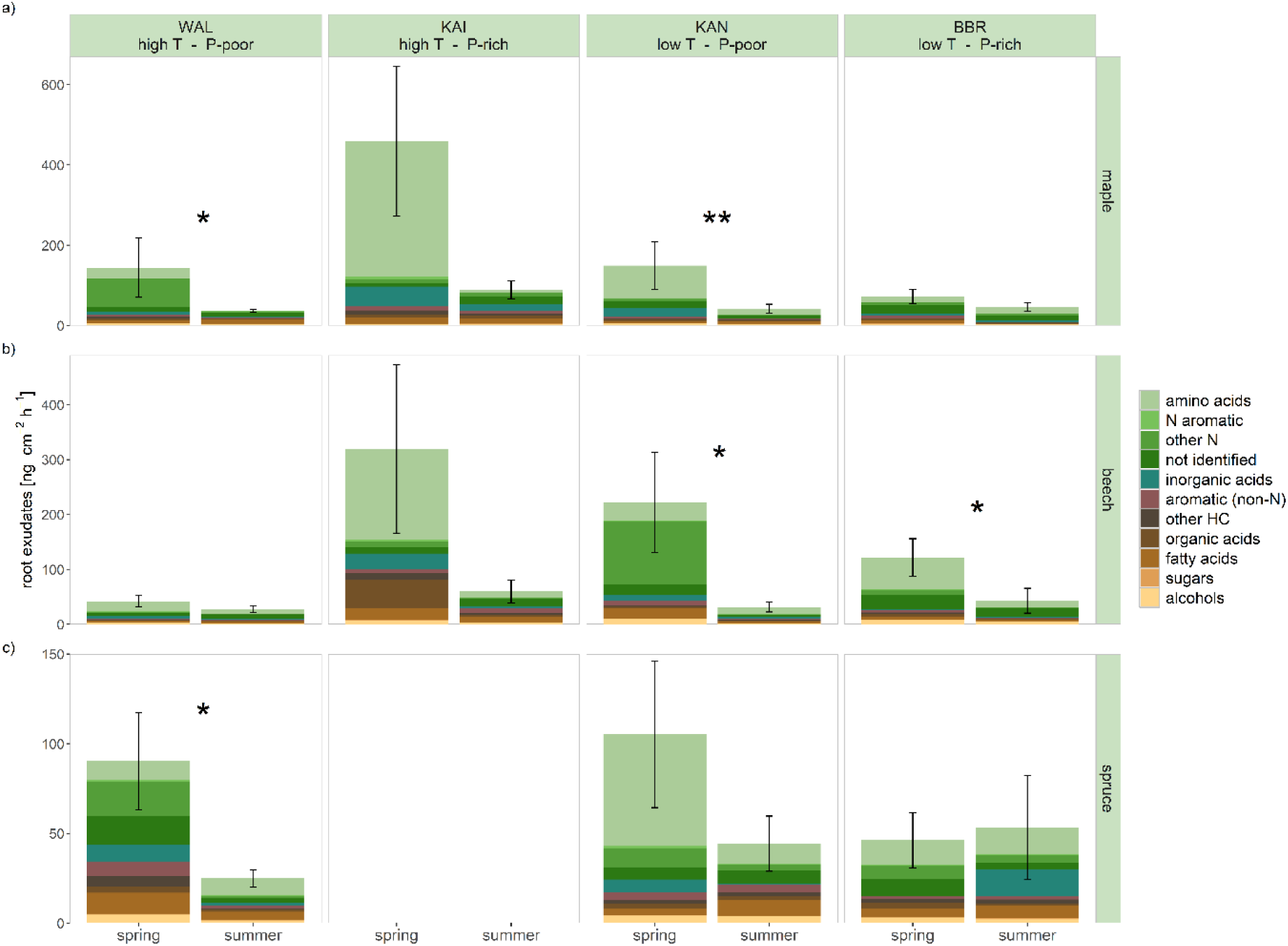
Root exudation in late spring (spring) and late summer (summer) for maple in a), beech in b), spruce in c) at the four study sites with different temperature (T) and soil phosphorus level (P). Colours code compound groups within the root exudates. Asterisks indicate significant differences (Wilcoxon signed-rank test) between late spring and late summer (p ≤ 0.01: **, p ≤ 0.05: *). Note the different scales for different species in a), b) and c).

### Seasonal variation

The sum of exudation in late spring was in general higher in all species across all sites than in late summer, except for spruce at BBR, where exudation was (non-significantly) higher in late summer (Figure 1). Maple and beech exuded more in late spring compared to late summer at all sites with significant differences at WAL and KAN for maple and at KAN and BBR in beech. Spruce also exuded more in late spring than in late summer, but only at the P-poor sites WAL (p ≤ 0.05) and KAN (p ≥ 0.05). At the low temperature P-rich site BBR, spruce trees exuded slightly less in late spring compared to late summer (Figure 1). The share of N-containing compounds also showed distinct seasonal patterns with a significantly higher share in late spring in all species. For examples, these compounds made up 68% in maple in late spring, but were reduced to 28% in late summer. In contrast, non-N-containing hydrocarbons, especially fatty acids and aromatics, were more dominant in late summer than in late spring in all species.

### Exudation in the forest floor and mineral soil

Figure 2 illustrates the different site- and species-specific exudation patterns in the forest floor and the mineral soil in late spring. A higher exudation in the mineral soil was found in spruce across all sites and in maple at the two warmer sites WAL and KAI (although only significant for spruce at BBR, p < 0.05). Contrarily, at KAN, the low-temperature, P-poor site with a comparatively thick forest floor, we observed a higher exudation in the forest floor in maple (not significant). No differences in exudation rates in forest floor and mineral soil were observed in maple at BBR and in beech at all sites except KAI (high temperature, P-rich). The share of all N-containing compounds in the mineral soil was higher (74% in maple to 56% in spruce), than in the forest floor (56% in maple to 43% in spruce). In contrast, the share of organic and fatty acids was higher in the forest floor compared to the mineral soil in all species. For instance, in beech, the share of organic acids in the forest floor was 12%, but only 2% of all exuded compounds were organic acids in the mineral soil. In general, variability in root exudation was high for all species at all sites. In late summer, similar exudation patterns for all species between forest floor and mineral soil were found, although differences were less pronounced (Figure S1).

**Figure 2:**
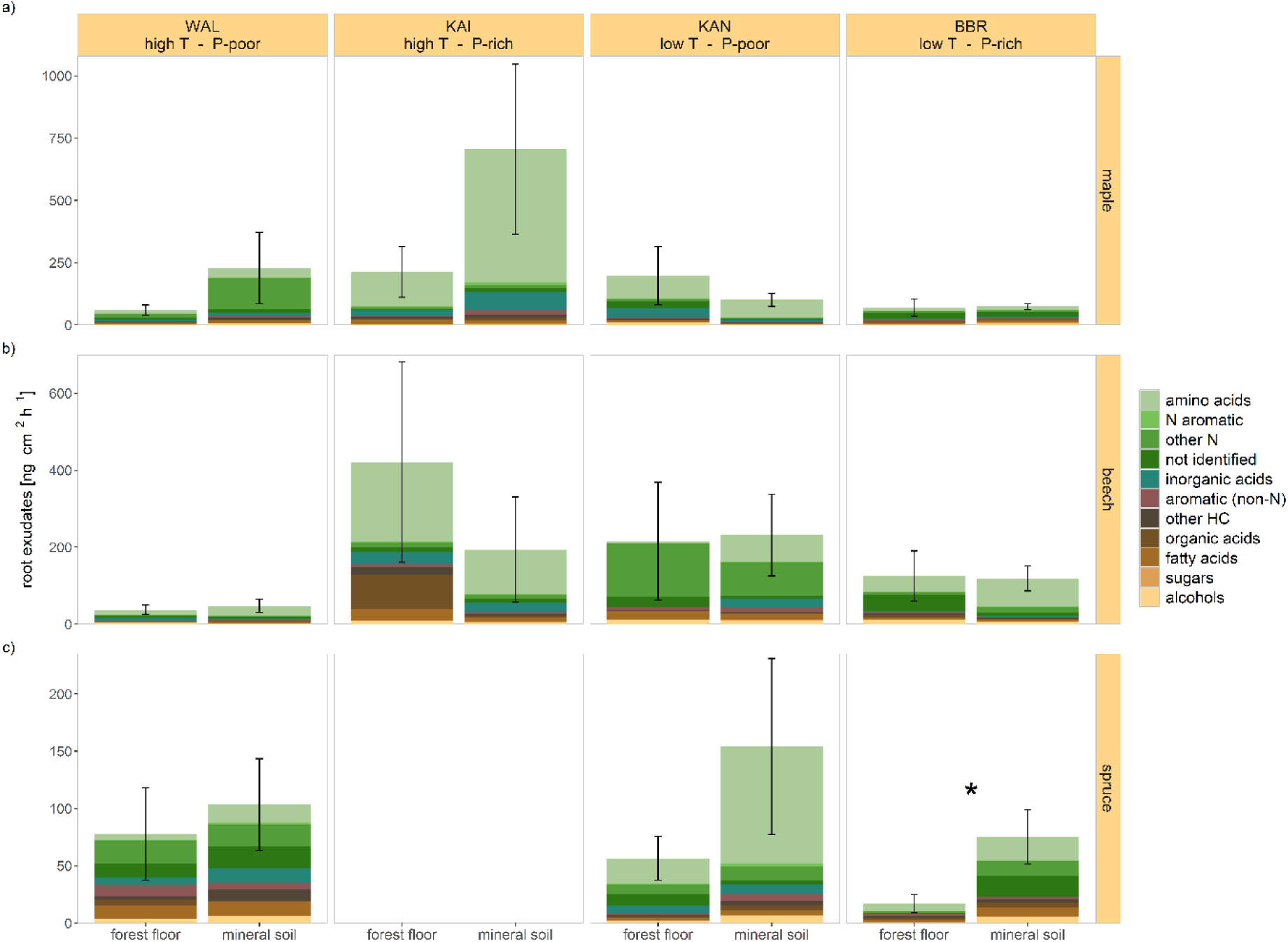
Root exudation in late spring in the forest floor and the mineral soil for maple in a), beech in b) and spruce in c) at the four study sites. Colours code compound groups within the exudates. Asterisks indicate significant differences between soil compartments (p ≤ 0.05). Note the different scales for different species.

### Exchangeable cations

Replicated soil elemental information of the four sites in the two soil compartments was used to examine relationships between root exudates and soil chemistry. Such relationships were most pronounced for exchangeable soil cations listed in Tables 4 - 6. Hereby, exchangeable base cations (Ca^2+^, K^+^, Mg^2+^ and Na^+^) and exchangeable non-base cations (Al^3+^, Fe^2+^ and Mn^2+^) led to distinct species- and compound-specific responses (exemplary illustrated in Figure 3). In general, Mn^2+^ was negatively correlated to the other exchangeable non-base cation leading to inversed relationships to root exudates compared to Al^3+^ and Fe^2+^. The representation in Figure 3 combines data from the forest floor and the mineral soil, since the observed trends were similar for both compartments. Graphs with separate regression lines for the two compartments can be found in the supplementary data (Figures S2 - S10).

**Figure 3:**
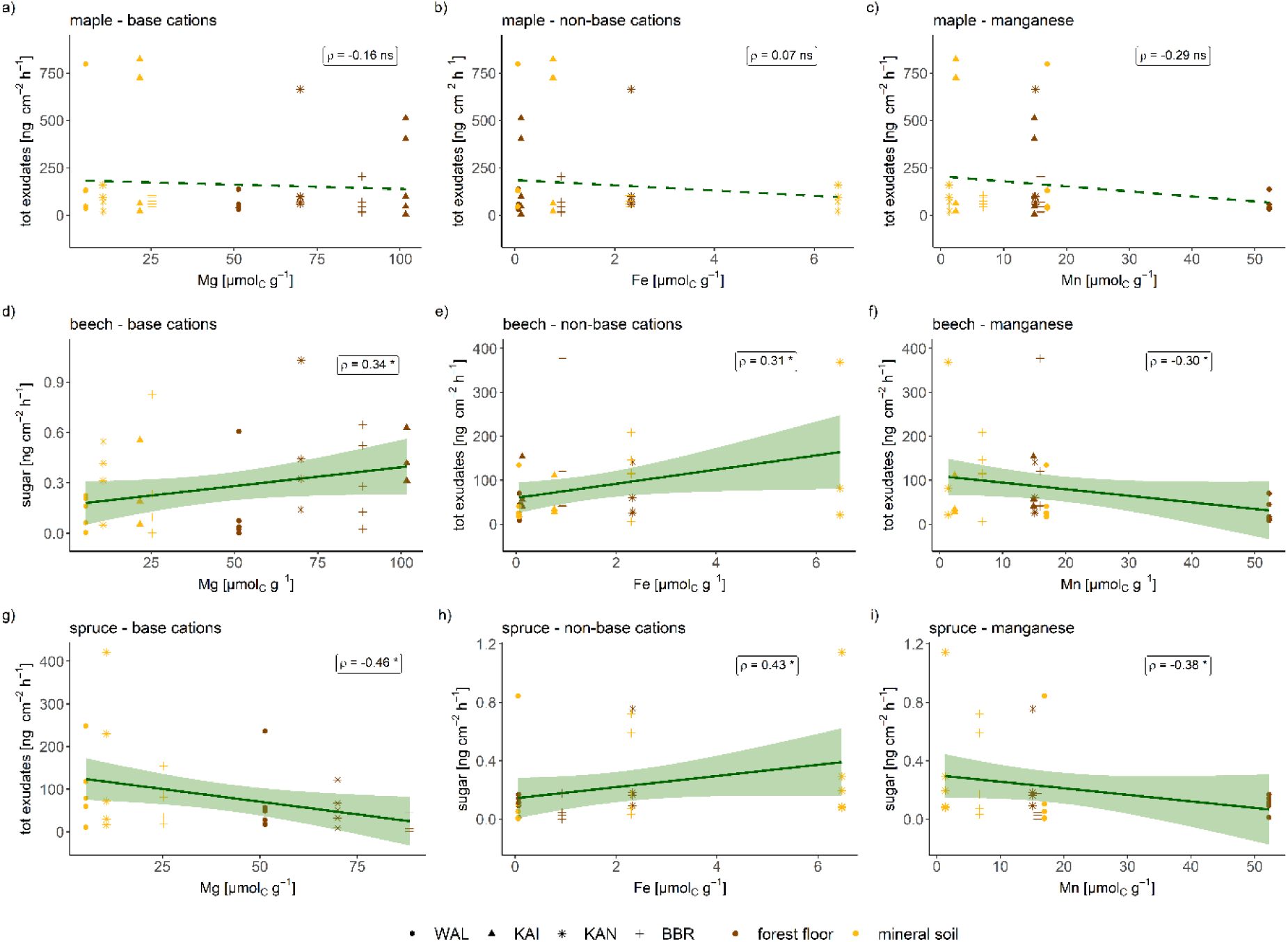
Exudate group in relation to exchangeable base cation concentration (a), d), g)), for exchangeable non-base cation concentration except Mn (b), e), h)) and exchangeable Mn concentration (c), f), i)) in maple (a) b), c)), in beech (d), e), f)) and in spruce (g), h), i)). Shapes indicate the four different sites and the colours the soil compartment. ρ indicates the Spearman’s rank correlation coefficient. Regression lines are illustrated by generalised linear smoothing and the shaded areas indicate the 95% confidence interval. Dashed regression lines are not significant. Note that variables and scales vary between panels.

In maple, the total exudation tended to decrease with increasing exchangeable base cation and to increase with increasing non-base cation concentration (except Mn^2+^). However, those tendencies were not statistically significant and some exudation compound groups, namely fatty acids and other N compounds, showed inverse patterns, i.e. a positive correlation with exchangeable base cations (Table 4).

**Table 4:** Correlation matrix of Spearman’s rank correlation coefficient ρ between compound groups in the exudates of maple in late spring and the concentration of exchangeable cations in the soil with base cations on the left and non-base cations on the right side of the matrix. Negative correlations are highlighted in shades of blue, positive ones in shades of red. Significant correlations are written in bold and marked with * for p ≤ 0.05 and ** for p ≤ 0.01.

| maple - $\rho$ | $\text{Ca}^{2+}$ | $\text{K}^{+}$ | $\text{Mg}^{2+}$ | $\text{Na}^{+}$ | $\text{Al}^{3+}$ | $\text{Fe}^{2+}$ | $\text{Mn}^{2+}$ |
| --- | --- | --- | --- | --- | --- | --- | --- |
| total exudation | -0.17 | -0.25 | -0.16 | -0.25 | 0.05 | 0.07 | -0.29 |
| amino acids | -0.25 | -0.26 | -0.28 | -0.30 | 0.13 | 0.08 | -0.23 |
| aromatic with N | -0.24 | -0.25 | -0.31 | -0.33 | 0.04 | -0.02 | -0.20 |
| other N compounds | 0.16 | 0.17 | 0.18 | 0.14 | -0.03 | 0.02 | 0.07 |
| other | -0.13 | -0.08 | -0.04 | -0.07 | 0.27 | 0.23 | -0.08 |
| inorganic acids | 0.07 | 0.00 | -0.02 | -0.06 | -0.14 | -0.07 | -0.12 |
| aromatic without N | -0.05 | -0.15 | -0.05 | -0.11 | -0.02 | -0.01 | -0.23 |
| other hydrocarbons | -0.22 | -0.23 | -0.31 | **-0.37\*** | -0.05 | -0.15 | -0.08 |
| organic acids | -0.13 | -0.19 | -0.10 | -0.16 | 0.10 | 0.12 | -0.24 |
| fatty acids | 0.17 | 0.11 | 0.12 | 0.07 | -0.27 | -0.27 | 0.09 |
| sugars | -0.08 | -0.04 | 0.14 | 0.15 | 0.41 | 0.40 | -0.15 |
| alcohols | -0.07 | -0.05 | 0.07 | 0.01 | 0.26 | 0.29 | -0.16 |

In beech, increasing exudation rates were observed for both, increasing exchangeable base and non-base cations (except Mn^2+^) across a wide range of exuded compound groups. Significant relationships could hereby be detected for Mg^2+^ and organic acids (Spearman’s rank correlation coefficient ρ = 0.27), including fatty acids (ρ = 0.29), sugars (ρ = 0.34) and other compounds (ρ = 0.33) for base cations. With respect to exchangeable non-base cations, Fe^2+^ and Mn^2+^ showed significant relationships with the total exudation rate (ρ = 0.31 for Fe^2+^, ρ = -0.30 for Mn^2+^), the exudation rate of other N compounds (ρ = 0.39 for Fe^2+^, ρ = -0.34 for Mn^2+^) and non-N aromatic compounds (ρ = 0.37 for Fe^2+^, ρ = -0.41 for Mn^2+^) in beech. While an increase with Fe^2+^ concentration was observed, there was a decrease with increasing Mn^2+^. Additionally, increasing Fe^2+^ went along with increasing exudation of alcohols (ρ = 0.33) and increasing Al^3+^ with increasing exudation of other N compounds (ρ = 0.34) (Table 5).

**Table 5:** Correlation matrix of Spearman’s rank correlation coefficient ρ between compound groups in the exudates of beech in late spring and the concentration of exchangeable cations in the soil with base cations on the left and non-base cations on the right side of the matrix. Negative correlations are highlighted in shades of blue, positive ones in shades of red. Significant correlations are written in bold and marked with * for p ≤ 0.05 and ** for p ≤ 0.01.

| beech - $\rho$ | $\text{Ca}^{2+}$ | $\text{K}^{+}$ | $\text{Mg}^{2+}$ | $\text{Na}^{+}$ | $\text{Al}^{3+}$ | $\text{Fe}^{2+}$ | $\text{Mn}^{2+}$ |
| --- | --- | --- | --- | --- | --- | --- | --- |
| total exudation | 0.06 | -0.03 | 0.23 | 0.14 | 0.14 | **0.31\*** | **-0.30\*** |
| amino acids | -0.08 | -0.07 | 0.03 | 0.03 | 0.11 | 0.07 | -0.04 |
| aromatic with N | -0.03 | -0.08 | 0.04 | 0.03 | -0.02 | 0.01 | -0.11 |
| other N compounds | -0.17 | -0.19 | -0.01 | -0.04 | **0.34\*** | **0.39\*** | **-0.34\*** |
| other | 0.26 | 0.25 | **0.33\*** | 0.26 | -0.02 | 0.15 | 0.03 |
| inorganic acids | 0.07 | 0.01 | -0.01 | 0.03 | -0.17 | -0.12 | -0.01 |
| aromatic without N | 0.04 | -0.07 | 0.23 | 0.10 | 0.18 | **0.37\*** | **-0.41\*\*** |
| other hydrocarbons | 0.02 | -0.08 | 0.26 | 0.20 | 0.15 | 0.26 | -0.20 |
| organic acids | 0.20 | 0.10 | **0.27\*** | 0.25 | -0.11 | 0.05 | -0.05 |
| fatty acids | 0.20 | 0.08 | **0.29\*** | 0.18 | -0.08 | 0.14 | -0.14 |
| sugars | 0.19 | 0.03 | **0.34\*** | 0.19 | -0.01 | 0.26 | -0.26 |
| alcohols | 0.06 | 0.04 | 0.22 | 0.16 | 0.19 | **0.33\*** | -0.17 |

In spruce, all exuded compounds decreased with increasing exchangeable base cations, while increasing non-base cations (except Mn^2+^) tended to increase exudation rates, but not in all exuded compounds. Generally, all analysed exchangeable base cations were significantly negatively correlated with the total exudation rates as well as the exudation rates of amino acids and alcohols (ρ ranging from -0.34 to -0.47). On the other hand, all analysed non-base cations (except Mn^2+^) showed a significant positive correlation with the exudation of sugars (ρ = 0.35 for Al^3+^, ρ = 0.43 for Fe^2+^, ρ = -0.38 for Mn^2+^) (Table 6).

**Table 6:** Correlation matrix of Spearman’s rank correlation coefficient ρ between compound groups in the exudates of spruce in late spring and the concentration of exchangeable cations in the soil with base cations on the left and non-base cations on the right side of the matrix. Negative correlations are highlighted in shades of blue, positive ones in shades of red. Significant correlations are written in bold and marked with * for p ≤ 0.05 and ** for p ≤ 0.01.

| spruce - $\rho$ | $\text{Ca}^{2+}$ | $\text{K}^{+}$ | $\text{Mg}^{2+}$ | $\text{Na}^{+}$ | $\text{Al}^{3+}$ | $\text{Fe}^{2+}$ | $\text{Mn}^{2+}$ |
| --- | --- | --- | --- | --- | --- | --- | --- |
| total exudation | **-0.34\*** | **-0.34\*** | **-0.46\*** | **-0.46\*** | 0.13 | 0.05 | -0.29 |
| amino acids | **-0.43\*** | **-0.43\*** | **-0.47\*\*** | **-0.47\*\*** | 0.21 | 0.12 | -0.34 |
| aromatic with N | -0.28 | -0.28 | **-0.43\*** | **-0.38\*** | -0.01 | -0.08 | -0.12 |
| other N compounds | -0.05 | -0.05 | -0.26 | -0.22 | -0.20 | -0.26 | 0.09 |
| other | -0.08 | -0.08 | -0.29 | -0.27 | -0.08 | -0.08 | -0.06 |
| inorganic acids | -0.13 | -0.13 | -0.20 | -0.20 | -0.08 | -0.06 | 0.00 |
| aromatic without N | -0.12 | -0.12 | -0.21 | -0.17 | 0.05 | 0.09 | -0.11 |
| other hydrocarbons | -0.14 | -0.14 | -0.27 | -0.23 | -0.08 | -0.20 | 0.00 |
| organic acids | -0.07 | -0.07 | -0.17 | -0.06 | 0.00 | 0.01 | -0.01 |
| fatty acids | -0.10 | -0.10 | **-0.37\*** | -0.29 | -0.24 | -0.33 | 0.10 |
| sugars | -0.19 | -0.19 | -0.07 | -0.16 | **0.35\*** | **0.43\*** | **-0.38\*** |
| alcohols | **-0.37\*** | **-0.37\*** | **-0.42\*** | **-0.39\*** | 0.23 | 0.13 | -0.33 |

## Discussion

The species-specific root exudation patterns observed in this study suggest that environmental conditions, such as season, MAT, organic or mineral soil compartments and nutrient availability are decisive factors for root exudation in temperate tree species. The three studied species showed distinct responses to these factors, stressing the importance of species-specific data. In our study, the AM associated maple showed a generally higher exudation rate and species responded differently to soil chemistry and soil compartment. Dynamics in root exudation were found even between roots of the same tree in different, but spatially close soil compartments, namely the forest floor and the mineral top soil (Figure 4). Some compound groups in the exudates, such as sugars and alcohols, reacted stronger to soil chemistry than others indicating not only quantitative, but also composition-specific alteration of root exudates in response to abiotic factors and site conditions.

**Figure 4:**
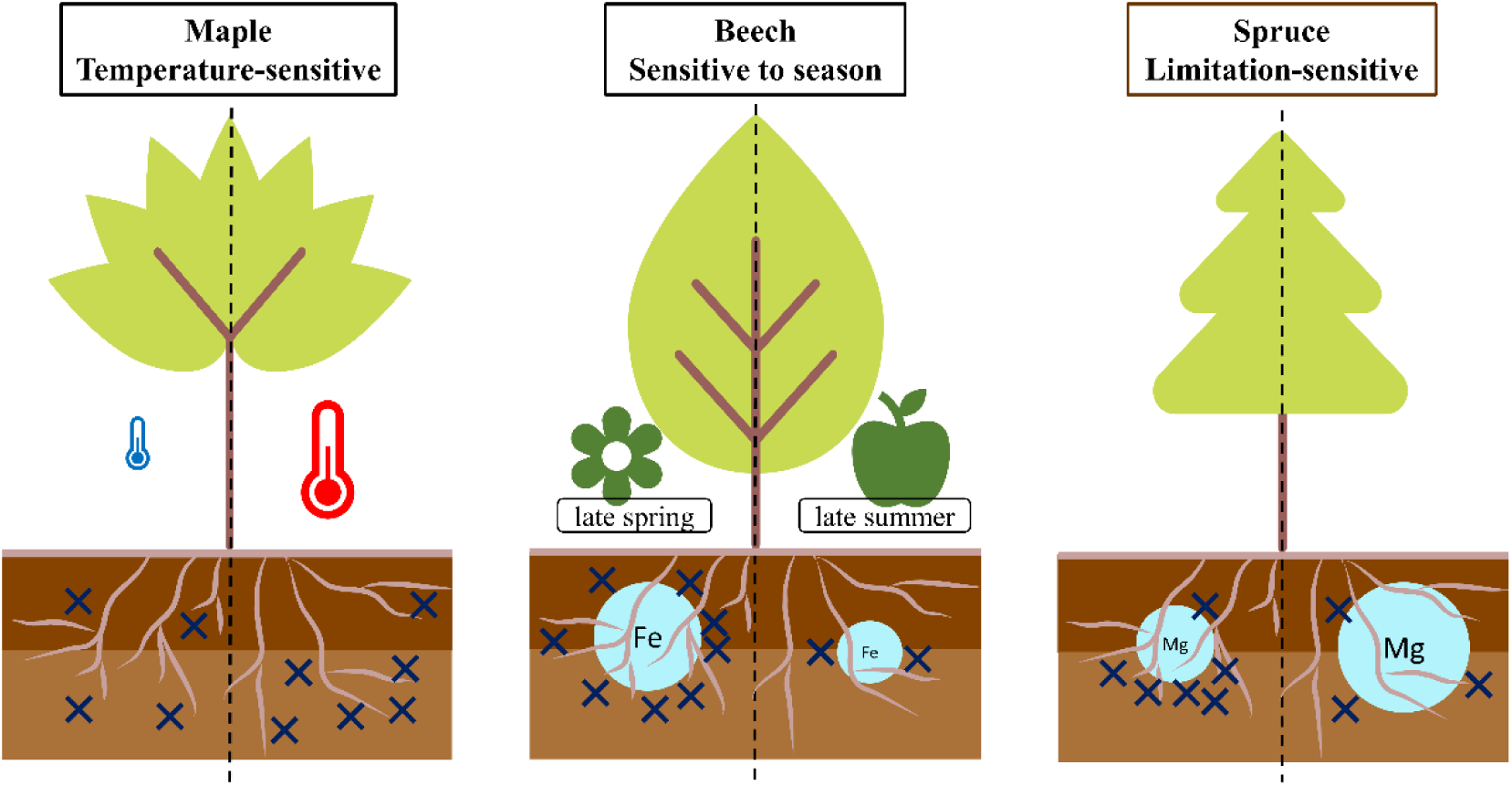
Conceptual representation of root exudation patterns (blue crosses) in maple, beech and spruce in the forest floor (dark brown) and the mineral soil (light brown) in response to temperature (coloured thermometers), season (flower: late spring, apple: late summer) and exchangeable soil cations (light blue circles).

### Species-specific root exudation patterns in temperate forests

The three studied tree species showed strong, site-dependent differences in exudation patterns. Such species differences were also observed in other studies; however, results are not consistent.

For example, it has been stated that EM associated species like beech and spruce generally exude more C than AM associated species like maple (Brzostek *et al*., 2015; Tückmantel *et al*., 2017). This is reasoned by the capability of EM-tree associations to produce extracellular enzymes, whereas AM-tree associations often lack this feature (Read & Perez-Moreno, 2003). Supporting this, Yin *et al*. (2014) observed that beech exuded a significantly higher amount of C than maple in three out of four sampling dates. However, in the present study, we observed a significantly higher exudation of soluble compounds in maple than in spruce, contradicting the above-mentioned findings. On average, maple individuals were thinner (and potentially younger) than the other species in this study, which could be a reason for an increased exudation. However, a correlation of root exudation to DBH in maple did not prove to be significant. Moreover, Tückmantel *et al*. (2017) brought forward that the reduced exudation in AM associated tree stands might be due to the higher bioavailability of nitrogen. This could explain the here observed patterns, since all sites in our study are beech-dominated and therefore also dominated by beech litter. Moreover, the higher exudation in maple could be a result of AM associations acquiring more nutrients, while beech and spruce are known to recycle more nutrients within the tree (Lang *et al*., 2016). Stronger N recycling could also explain the tendency of beech and spruce to exude a lower share of N-containing compounds compared to maple. Besides, it has been argued that the decreased meristematic activity under P-limiting conditions would lead to a decreased exudation at P-poor sites (Canarini *et al*., 2019). However, other studies observed the inverse (Jones *et al*., 2004; Jones *et al*., 2009; Vives-Peris *et al*., 2020) or no relationship at all in beech (Leuschner *et al*., 2022). Supporting the latter finding, we found beech and maple to exude similar amounts at P-rich and P-poor sites, while spruce exuded more at P-poor sites (even though not significantly). However, it needs to be pointed out that these discrepancies could potentially also arise from varying methods of analysis. While it is rather new to apply a compounds-specific approach to root exudates of temperate forest trees, many studies so far focused on total exuded carbon. It also becomes clear that the site conditions evaluated here could not explain the site effect on root exudation completely, which suggests complex interaction and additional factors affecting root exudation. To elucidate these, further research is needed.

### Seasonal dynamics in root exudation in temperate tree species

Although there is a seasonal course of photosynthetic C assimilation in temperate forests (Carrara *et al*., 2004; Tyrrell *et al*., 2012), seasonal patterns in root exudation have not been found consistently (Phillips *et al*., 2008; Tückmantel *et al*., 2017).

In this study, however, we detected a difference between late spring and late summer with generally higher exudation in late spring, when C assimilation is also higher. Similarly, higher exudation in summer compared to winter has been observed before (Chen *et al*., 2023). Further, in an early stage of the vegetation period, tree growth and therefore nutrient uptake is stronger compared to the late stage of the vegetation period with higher uptake of P and N in spring (May) compared to autumn (September-October) in beech at three sites including one studied here (Likulunga *et al*., 2022). Enhanced nutrient uptake could therefore explain the increased exudation in late spring. This is supported by the higher share of N-containing compounds in late spring, which accelerates the biomass decomposition through a lower C/N ratio in the exudates (Meier *et al*., 2017). Adding to this, the net primary production of temperate forests is higher in May/June than in September, but respiration is still high in September (Carrara *et al*., 2004). This suggests that the trees are still active in September but with a reduced nutrient demand. Next to the seasonal variation in nutrient uptake, there was a difference in drought index between the two seasons in 2023. The standardised precipitation evaporation index (SPEI) (Vicente-Serrano *et al*., 2010) was lower in late summer than in late spring at all sites. The SPEI (calculated based on the six months before sampling) changed from mild drought in late spring to moderate drought in late summer at the sites WAL, KAI and KAN. At BBR, no indication of drought was evident in late spring, but a mild drought was shown in late summer by the index. The reduced C assimilation under drought (Bréda *et al*., 2006) could therefore also be a reason for the smaller exudation rates in late summer. This is supported by Dannenmann *et al*. (2009) showing reduced exudation rates under drought. However, a range of studies found the opposite effect (Jakoby *et al*., 2020; Meier *et al*., 2020; Brunn *et al*., 2022; Leuschner *et al*., 2022). This advocates for more drought-related root exudation studies in temperate tree species (Möller *et al*., 2024).

### Root exudation in the forest floor and the mineral soil

In this study, we found a comparatively low exudation in maple in the forest floor at the warm sites WAL and KAI, which have a thinner forest floor and therefore a lower share of organically-bound nutrients, while the exudation at the colder P-poor site KAN with a thick forest floor is higher in the forest floor compared to the mineral soil. The quantity of plant nutrients present in the organic or inorganic form was found to influence root exudation in various studies. Hereby, a positive relationship between organically bound N in the soil and root exudation C was found (Rohrbacher & St-Arnaud, 2016; Tückmantel *et al*., 2017; Vives-Peris *et al*., 2020). Adding to this, it has been stated that the often-observed increased exudation under N- and P-limiting conditions (Jones *et al*., 2004; Jones *et al*., 2009; Vives-Peris *et al*., 2020) only holds when N is present in the organic form (Meier *et al*., 2020). This could be a mechanism to increase the carbon use efficiency by avoiding the production of compounds for priming effects when it is inefficient, because microbial activity cannot increase the bioavailability of N if not present as organic N. This could explain our observed pattern in maple. Additionally, higher temperatures at WAL and KAI also most likely lead to a generally higher mineralisation rate, which could decrease the need of exudates for mineralisation. This pattern is not present in beech and spruce, which might be due to their above-mentioned higher nutrient recycling capacity compared to maple, which belongs to the nutrient acquiring type (Lang *et al*., 2016). Those different nutrient economies are represented in the litter quality being low in nutrient-recycling types, especially in spruce having a high C/N ratio in the litter (Vesterdal *et al*., 2008) and therefore in the forest floor. In contrast, the C/N ratio in the SOM in the mineral soil seems to be similar for the three species (Vesterdal *et al*., 2013). This could explain the increased exudation of spruce in the mineral soil, especially at the colder sites, where decomposition is slow. Even though many of the differences between soil compartments are not significant, they indicate that trees can adjust their root exudates on a very small scale, as between different soil compartments.

### Root exudation in temperate tree species in response to soil chemistry

The capacity of dynamic adjustment is also represented in the significant relationships detected between root exudates and soil elements (Figure 3). We mainly identified significant relationships for root exudates versus exchangeable cations, representing the fact that the exchangeable form of elements is most important for temperate tree species. However, it remains unclear whether the abundance of exchangeable elements exerts an effect on the root exudates or vice versa. Malate, pyruvate and citrate have been identified to be helpful to withstand Al toxicity (Pantigoso *et al*., 2021). Previous studies have already found indications of Al^3+^ complexation through root exudates to reduce phytotoxic effects (Heim *et al*., 2001; Collignon *et al*., 2012). This could explain the positive relationship between the exudation of hydrocarbons and exchangeable non-base cations in the soil in beech and spruce in our study. In our case, other N hydrocarbons (for beech) and sugars (for spruce) were involved, which could be a species-specific reaction. The same could hold for beech, which also increased exudation with increasing non-base cation concentration. In beech, however, Fe^2+^ seemed to have a stronger effect than Al^3+^, which could also be a species-specific reaction.

The relationship between base cations on root exudation is less studied. Interestingly, beech and spruce showed inverse patterns in this regard. The increased rate of total exudation, amino acid and alcohol exudation in spruce under decreased exchangeable base cation concentration in the soil could increase the mobility of those elements by desorbing them from e.g. clay minerals (alcohols and amino acids) and decreasing the C/N ratio in the soil (amino acids) which accelerates the SOM degradation through microorganisms (Meier *et al*., 2017). The latter mechanism is especially relevant for spruce because of the high C/N ratio in its litter (Vesterdal *et al*., 2013). In contrast, N-containing compounds in the exudates of beech were not related to the abundance of exchangeable base cations. Instead, we found the exudation of organic acids, fatty acids and sugars to correlate positively with the exchangeable base cation Mg^2+^. The observed pattern could be a hint for higher Mg^2+^ concentrations being a consequence and not a cause of exudation by beech. This mechanism has been observed for organic acid exudation in *Cryptomeria japonica* (Ohta & Hiura, 2016). However, this effect could not be seen in the other species and needs more research to be fully understood. Further compound targeted studies could reveal the specific role of beech in this case. Nonetheless, it is a further indicator for pronounced species differences and shows that not only the rate of exudation, but also its composition is dynamically adjusted to abiotic factors.

## Conclusion

While spruce shows a tendency to exude more in the mineral soil and reacted sensitively to nutrient availability with increased exudation under limiting conditions of P and base cations (limitation-sensitive exudation), maple seems to response stronger to temperature with higher exudation rates at warm sites, especially in the mineral soil (temperature-sensitive exudation). The exudation of beech was related to the concentration of non-base cations with higher concentration inducing higher exudation, which might be a protection mechanism against phytotoxicity, but was not related to soil compartment nor temperature. However, beech showed the highest decrease in exudation between late spring and late summer (exudation sensitive to season) (Figure 4).

This strongly indicates species-specific adaptations and dynamics of root exudation to season, soil compartment and soil chemistry. Interestingly, roots from the same trees showed differences in exudation in the two soil compartments. This suggests that an active response in root exudation to specific conditions in the forest floor, with lower risk of Al toxicity and higher share of organically bound nutrients is possible. The dynamics in root exudation was also reflected in the seasonal patterns and adjustments to the abundance of exchangeable soil cations.

The results of this study show that root exudation reacts dynamically to environmental parameters. Therefore, gaining knowledge about root exudates, especially in field environments, is particularly challenging. Analysing the composition next to the rate of exudates allows more detailed insights in root exudation patterns, drivers and functions. Compound analyses might give a clue to so far often contrasting findings. For instance, in this study a strong response of the sugar exudation in spruce to the concentration of non-base cation was found, which was not significantly reflected in total exudation rates.

## Author contributions

CW, JK, MW and SH designed the study. MW conducted the field and lab work. JK assisted during sample processing and analysis. MW drafted the manuscript with major contributions of SH. CW and SH helped with data analysis and interpretation. JN, JP and FL contributed site and soil chemistry data. All authors critically reviewed the manuscript and contributed substantially.

## Acknowledgements

This study was carried out in the framework of the Research Unit 5315 “Forest Floor: Functioning, Dynamics and Vulnerability in a Changing World” funded by the DFG (project no.: WE2681/13-1). We thank all research unit members and the working group of the Ecosystem Physiology, especially Monika Eiblmeier and Alexandra Paul for lab assistance and analyses and the students Marie Rein and Lee Deutsch for their help in the field and lab.

## Data Availability

The data that support the findings of this study are available from the corresponding author upon reasonable request.

## References

Beguería, S., Vicente-Serrano, S.M. & Angulo-Martínez, M. (2010) A Multiscalar Global Drought Dataset: The SPEIbase: A New Gridded Product for the Analysis of Drought Variability and Impacts. Bulletin of the American Meteorological Society, 91(10), 1351–1356. Available from: 10.1175/2010BAMS2988.1.

Bréda, N., Huc, R., Granier, A. & Dreyer, E. (2006) Temperate forest trees and stands under severe drought: a review of ecophysiological responses, adaptation processes and long-term consequences. Annals of Forest Science, 63(6), 625–644. Available from: 10.1051/forest:2006042.

Brunn, M., Hafner, B.D., Zwetsloot, M.J., Weikl, F., Pritsch, K. & Hikino, K. et al. (2022) Carbon allocation to root exudates is maintained in mature temperate tree species under drought. New Phytologist, 235(3), 965–977. Available from: 10.1111/nph.18157.

Brzostek, E.R., Dragoni, D., Brown, Z.A. & Phillips, R.P. (2015) Mycorrhizal type determines the magnitude and direction of root-induced changes in decomposition in a temperate forest. New Phytologist, 206(4), 1274–1282. Available from: 10.1111/nph.13303.

Canarini, A., Kaiser, C., Merchant, A., Richter, A. & Wanek, W. (2019) Root Exudation of Primary Metabolites: Mechanisms and Their Roles in Plant Responses to Environmental Stimuli. Frontiers in Plant Science, 10, 157. Available from: 10.3389/fpls.2019.00157.

Carrara, A., Janssens, I.A., Curiel Yuste, J. & Ceulemans, R. (2004) Seasonal changes in photosynthesis, respiration and NEE of a mixed temperate forest. Agricultural and Forest Meteorology, 126(1-2), 15–31. Available from: 10.1016/j.agrformet.2004.05.002.

Chen, M., Yao, X., Cheng, H., Fan, A., Lin, R. & Wang, X. et al. (2023) Changes in Chinese fir plantations root exudation strategies seasonally and as tree age. Forest Ecology and Management, 545, 121239. Available from: 10.1016/j.foreco.2023.121239.

Collignon, C., Boudot, J.-P. & Turpault, M.-P. (2012) Time change of aluminium toxicity in the acid bulk soil and the rhizosphere in Norway spruce (Picea abies (L.) Karst.) and beech (Fagus sylvatica L.) stands. Plant and Soil, 357(1-2), 259–274. Available from: 10.1007/s11104-012-1154-2.

Dannenmann, M., Simon, J., Gasche, R., Holst, J., Naumann, P.S. & Kögel-Knabner, I. et al. (2009) Tree girdling provides insight on the role of labile carbon in nitrogen partitioning between soil microorganisms and adult European beech. Soil Biology and Biochemistry, 41(8), 1622–1631. Available from: 10.1016/j.soilbio.2009.04.024.

DeForest, J.L. & Snell, R.S. (2020) Tree growth response to shifting soil nutrient economy depends on mycorrhizal associations. New Phytologist, 225(6), 2557–2566. Available from: 10.1111/nph.16299.

Finzi, A.C., Abramoff, R.Z., Spiller, K.S., Brzostek, E.R., Darby, B.A. & Kramer, M.A. et al. (2015) Rhizosphere processes are quantitatively important components of terrestrial carbon and nutrient cycles. Global Change Biology, 21(5), 2082–2094. Available from: 10.1111/gcb.12816.

Heim, A., Brunner, I., Frey, B., Frossard, E. & Luster, J. (2001) Root exudation, organic acids, and element distribution in roots of Norway spruce seedlings treated with aluminum in hydroponics. Journal of Plant Nutrition and Soil Science, 164(5), 519. Available from: 10.1002/1522-2624(200110)164:5<519::AID-JPLN519>3.0.CO;2-Y.

Huang, X.-F., Chaparro, J.M., Reardon, K.F., Zhang, R., Shen, Q. & Vivanco, J.M. (2014) Rhizosphere interactions: root exudates, microbes, and microbial communities. Botany, 92(4), 267–275. Available from: 10.1139/cjb-2013-0225.

IUSS Working Group WRB (2022) World Reference Base for Soil Resources: International soil classification system for naming soils and creating legends for soil maps, 4th Edition.

Jakoby, G., Rog, I., Megidish, S. & Klein, T. (2020) Enhanced root exudation of mature broadleaf and conifer trees in a Mediterranean forest during the dry season. Tree Physiology, 40(11), 1595–1605. Available from: 10.1093/treephys/tpaa092.

Jones, D.L., Hodge, A. & Kuzyakov, Y. (2004) Plant and mycorrhizal regulation of rhizodeposition. New Phytologist, 163(3), 459–480. Available from: 10.1111/j.1469-8137.2004.01130.x.

Jones, D.L., Nguyen, C. & Finlay, R.D. (2009) Carbon flow in the rhizosphere: carbon trading at the soil–root interface. Plant and Soil, 321(1-2), 5–33. Available from: 10.1007/s11104-009-9925-0.

Kreuzwieser, J., Hauberg, J., Howell, K.A., Carroll, A., Rennenberg, H. & Millar, A.H. et al. (2009) Differential response of gray poplar leaves and roots underpins stress adaptation during hypoxia. Plant Physiology, 149(1), 461–473. Available from: 10.1104/pp.108.125989.

Lang, F., Bauhus, J., Frossard, E., George, E., Kaiser, K. & Kaupenjohann, M. et al. (2016) Phosphorus in forest ecosystems: New insights from an ecosystem nutrition perspective. Journal of Plant Nutrition and Soil Science, 179(2), 129–135. Available from: 10.1002/jpln.201500541.

Lang, F., Krüger, J., Amelung, W., Willbold, S., Frossard, E. & Bünemann, E.K. et al. (2017) Soil phosphorus supply controls P nutrition strategies of beech forest ecosystems in Central Europe. Biogeochemistry, 136(1), 5–29. Available from: 10.1007/s10533-017-0375-0.

Leuschner, C., Tückmantel, T. & Meier, I.C. (2022) Temperature effects on root exudation in mature beech (Fagus sylvatica L.) forests along an elevational gradient. Plant and Soil, 481(1-2), 147–163. Available from: 10.1007/s11104-022-05629-5.

Likulunga, L.E., Clausing, S., Krüger, J., Lang, F. & Polle, A. (2022) Fine root biomass of European beech trees in different soil layers show different responses to season, climate, and soil nutrients. Frontiers in Forests and Global Change, 5, 955327. Available from: 10.3389/ffgc.2022.955327.

Massalha, H., Korenblum, E., Tholl, D. & Aharoni, A. (2017) Small molecules below-ground: the role of specialized metabolites in the rhizosphere. The Plant Journal : for Cell and Molecular Biology, 90(4), 788–807. Available from: 10.1111/tpj.13543.

Maurer, D., Malique, F., Alfarraj, S., Albasher, G., Horn, M.A. & Butterbach-Bahl, K. et al. (2021) Interactive regulation of root exudation and rhizosphere denitrification by plant metabolite content and soil properties. Plant and Soil, 467(1-2), 107–127. Available from: 10.1007/s11104-021-05069-7.

Meier, I.C., Finzi, A.C. & Phillips, R.P. (2017) Root exudates increase N availability by stimulating microbial turnover of fast-cycling N pools. Soil Biology and Biochemistry, 106, 119–128. Available from: 10.1016/j.soilbio.2016.12.004.

Meier, I.C., Tückmantel, T., Heitkötter, J., Müller, K., Preusser, S. & Wrobel, T.J. et al. (2020) Root exudation of mature beech forests across a nutrient availability gradient: the role of root morphology and fungal activity. New Phytologist, 226(2), 583–594. Available from: 10.1111/nph.16389.

Möller, K., Ritter, A., Stobinsky, P.J., Jensen, K., Meier, I.C. & Subrahmaniam, H.J. (2024) Targeting the untargeted: Uncovering the chemical complexity of root exudates. BioRxiv, 2024.09.17.613458. Available from: 10.1101/2024.09.17.613458.

Neumann, G., Bott, S., Ohler, M.A., Mock, H.-P., Lippmann, R. & Grosch, R. et al. (2014) Root exudation and root development of lettuce (Lactuca sativa L. cv. Tizian) as affected by different soils. Frontiers in Microbiology, 5. Available from: 10.3389/fmicb.2014.00002.

Ohta, T. & Hiura, T. (2016) Root exudation of low-molecular-mass-organic acids by six tree species alters the dynamics of calcium and magnesium in soil. Canadian Journal of Soil Science, 96(2), 199–206. Available from: 10.1139/cjss-2015-0063.

Pantigoso, H.A., He, Y., DiLegge, M.J. & Vivanco, J.M. (2021) Methods for Root Exudate Collection and Analysis. In: Carvalhais, L.C. & Dennis, P.G. (Eds.) The Plant Microbiome: Methods and Protocols. Springer US; Imprint: Humana: New York, NY.

Phillips, R.P., Brzostek, E. & Midgley, M.G. (2013) The mycorrhizal-associated nutrient economy: a new framework for predicting carbon-nutrient couplings in temperate forests. New Phytologist, 199(1), 41–51. Available from: 10.1111/nph.12221.

Phillips, R.P., Erlitz, Y., Bier, R. & Bernhardt, E.S. (2008) New approach for capturing soluble root exudates in forest soils. Functional Ecology, 22(6), 990–999. Available from: 10.1111/j.1365-2435.2008.01495.x.

Phillips, R.P., Finzi, A.C. & Bernhardt, E.S. (2011) Enhanced root exudation induces microbial feedbacks to N cycling in a pine forest under long-term CO2 fumigation. Ecology Letters, 14(2), 187–194. Available from: 10.1111/j.1461-0248.2010.01570.x.

Prescott, C.E. & Grayston, S.J. (2013) Tree species influence on microbial communities in litter and soil: Current knowledge and research needs. Forest Ecology and Management, 309, 19–27. Available from: 10.1016/j.foreco.2013.02.034.

Read, D.J. & Perez-Moreno, J. (2003) Mycorrhizas and nutrient cycling in ecosystems - a journey towards relevance? New Phytologist, 157(3), 475–492. Available from: 10.1046/j.1469-8137.2003.00704.x.

Rohrbacher, F. & St-Arnaud, M. (2016) Root Exudation: The Ecological Driver of Hydrocarbon Rhizoremediation. Agronomy, 6(1), 19. Available from: 10.3390/agronomy6010019.

Ruf, A., Kuzyakov, Y. & Lopatovskaya, O. (2006) Carbon fluxes in soil food webs of increasing complexity revealed by 14C labelling and 13C natural abundance. Soil Biology and Biochemistry, 38(8), 2390–2400. Available from: 10.1016/j.soilbio.2006.03.008.

Tückmantel, T., Leuschner, C., Preusser, S., Kandeler, E., Angst, G. & Mueller, C.W. et al. (2017) Root exudation patterns in a beech forest: Dependence on soil depth, root morphology, and environment. Soil Biology and Biochemistry, 107, 188–197. Available from: 10.1016/j.soilbio.2017.01.006.

Tyrrell, M.L., Ross, J. & Kelty, M. (2012) Carbon Dynamics in the Temperate Forest. In: Ashton, M.S. (Ed.) Managing forest carbon in a changing climate. Springer: Dordrecht, New York, pp. 77–107.

Vesterdal, L., Clarke, N., Sigurdsson, B.D. & Gundersen, P. (2013) Do tree species influence soil carbon stocks in temperate and boreal forests? Forest Ecology and Management, 309, 4–18. Available from: 10.1016/j.foreco.2013.01.017.

Vesterdal, L., Schmidt, I.K., Callesen, I., Nilsson, L.O. & Gundersen, P. (2008) Carbon and nitrogen in forest floor and mineral soil under six common European tree species. Forest Ecology and Management, 255(1), 35–48. Available from: 10.1016/j.foreco.2007.08.015.

Vicente-Serrano, S.M., Beguería, S. & López-Moreno, J.I. (2010) A Multiscalar Drought Index Sensitive to Global Warming: The Standardized Precipitation Evapotranspiration Index. Journal of Climate, 23(7), 1696–1718. Available from: 10.1175/2009JCLI2909.1.

Vives-Peris, V., Ollas, C. de, Gómez-Cadenas, A. & Pérez-Clemente, R.M. (2020) Root exudates: from plant to rhizosphere and beyond. Plant Cell Reports, 39(1), 3–17. Available from: 10.1007/s00299-019-02447-5.

Vranova, V., Rejsek, K., Skene, K.R., Janous, D. & Formanek, P. (2013) Methods of collection of plant root exudates in relation to plant metabolism and purpose: A review. Journal of Plant Nutrition and Soil Science, 176(2), 175–199. Available from: 10.1002/jpln.201000360.

Wachendorf, C., Frank, T., Broll, G., Beylich, A. & Milbert, G. (2023) A Concept for a Consolidated Humus Form Description—An Updated Version of German Humus Form Systematics. International Journal of Plant Biology, 14(3), 658–686. Available from: 10.3390/ijpb14030050.

Williams, A., Langridge, H., Straathof, A.L., Muhamadali, H., Hollywood, K.A. & Goodacre, R. et al. (2022) Root functional traits explain root exudation rate and composition across a range of grassland species. Journal of Ecology, 110(1), 21–33. Available from: 10.1111/1365-2745.13630.

Yin, H., Wheeler, E. & Phillips, R.P. (2014) Root-induced changes in nutrient cycling in forests depend on exudation rates. Soil Biology and Biochemistry, 78, 213–221. Available from: 10.1016/j.soilbio.2014.07.022.

