## Supplementary data for "Spatio-temporal plasticity of root exudation in three temperate tree species: effects of season, site and soil characteristics"

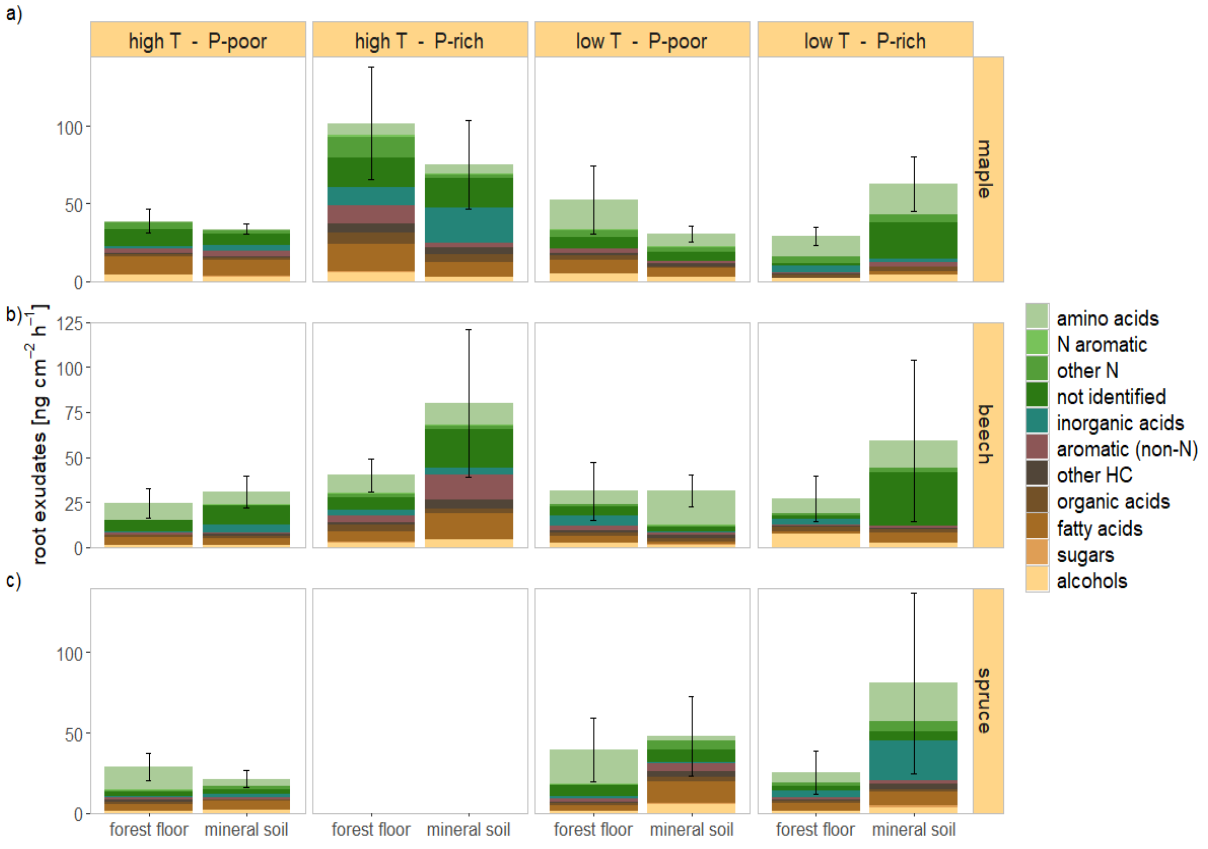


***Figure S1***: Root exudation in late summer in the forest floor and the mineral soil for maple in a), beech in b) and spruce in c) at the four study sites. Colours code compound groups within the exudates. Note the different scales for different species.

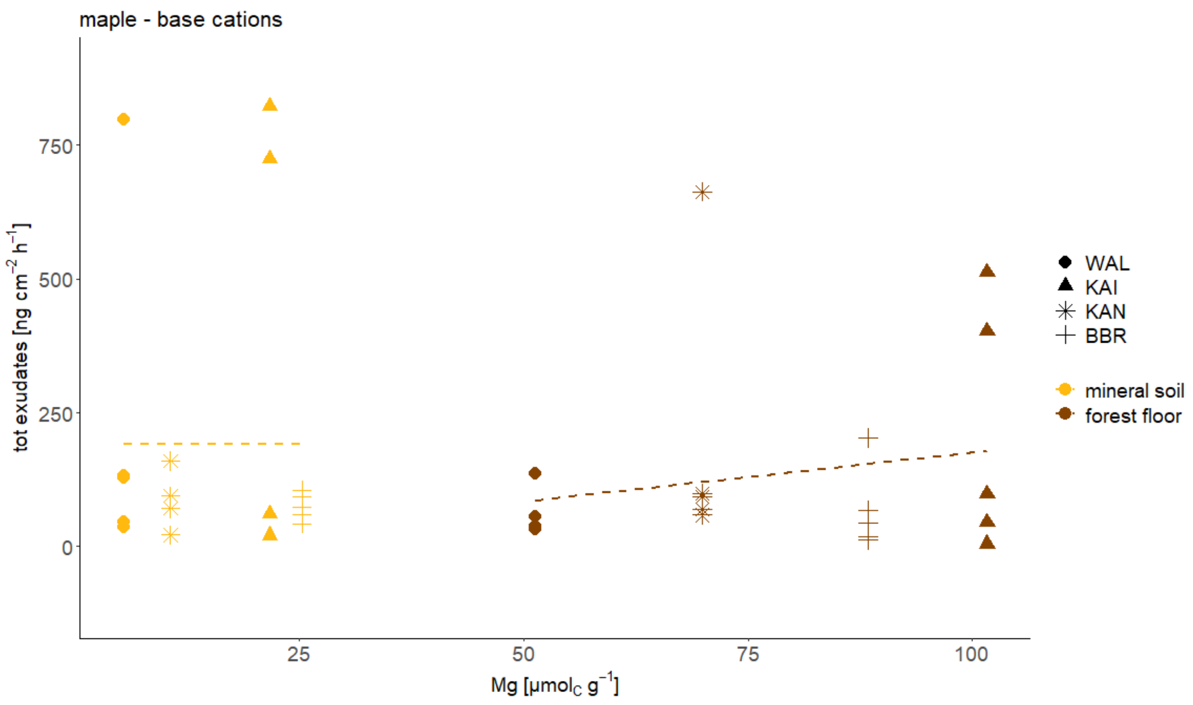


***Figure S2***: Total exudates in relation to exchangeable Mg concentration for maple. Shapes indicate the four different sites and the colours the horizon. Regression lines are illustrated by generalized linear smoothing.

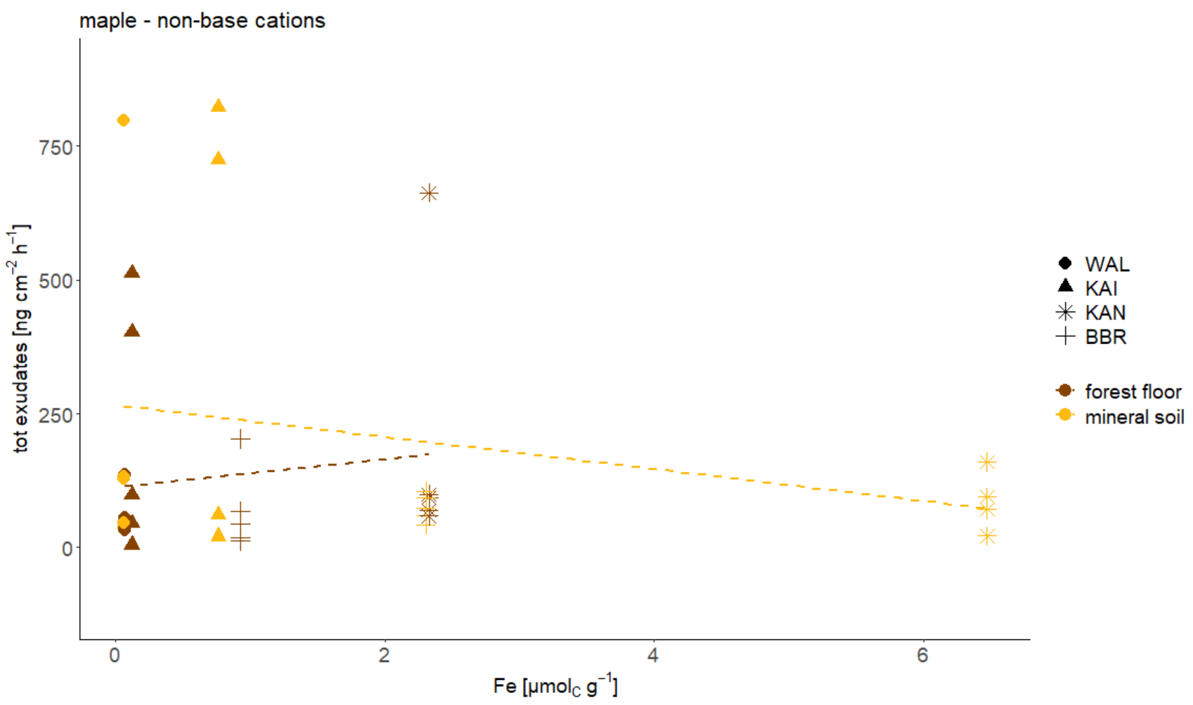


***Figure S3***: Total exudates in relation to exchangeable Fe concentration for maple. Shapes indicate the four different sites and the colours the horizon. Regression lines are illustrated by generalized linear smoothing.

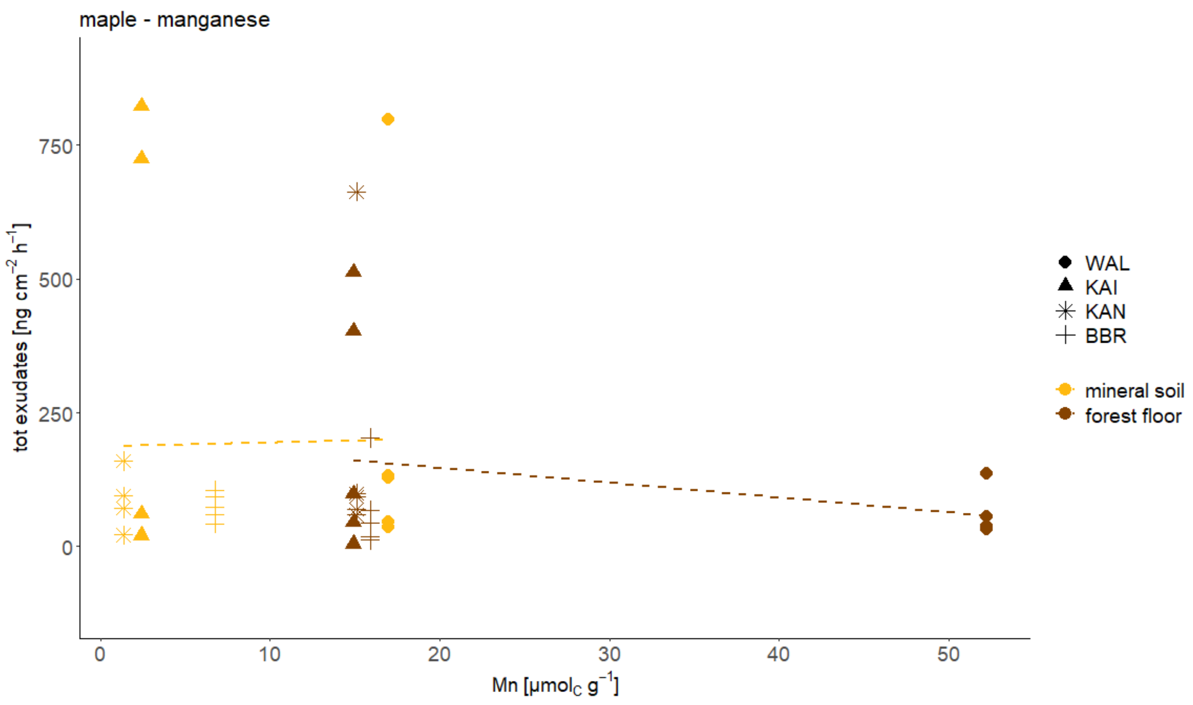


***Figure S4***: Total exudates in relation to exchangeable Mn concentration for maple. Shapes indicate the four different sites and the colours the horizon. Regression lines are illustrated by generalized linear smoothing.

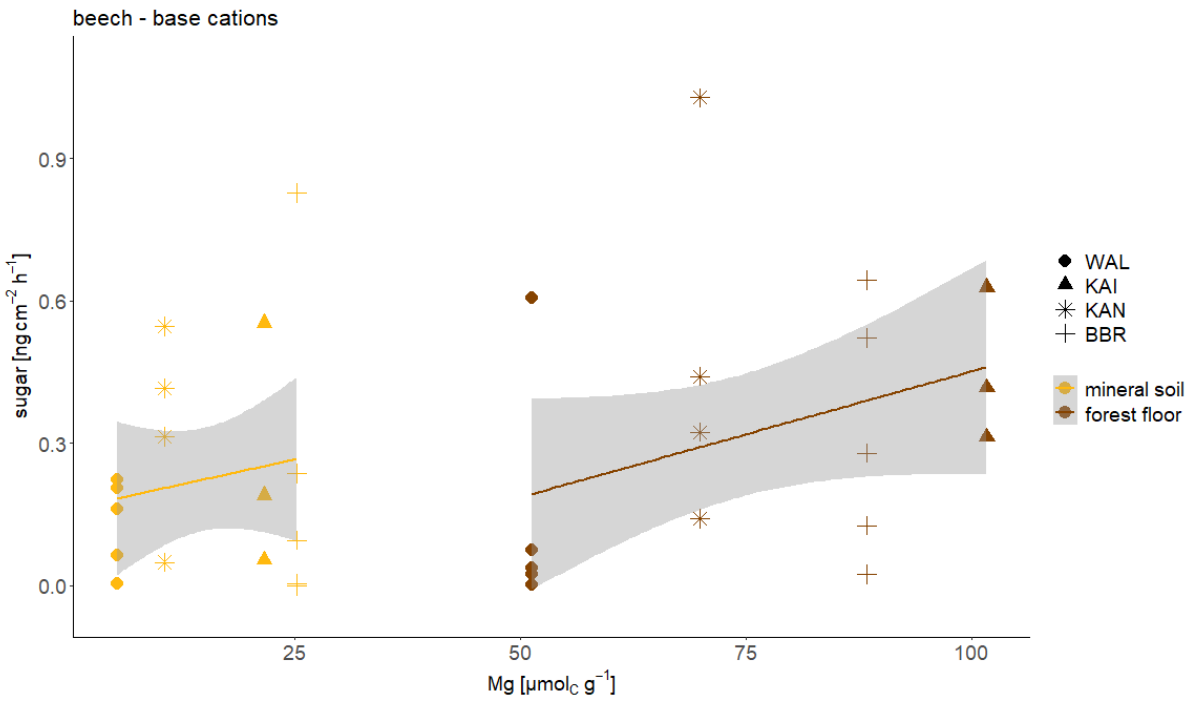


***Figure S5***: Sugar exudation in relation to exchangeable Mg concentration for beech. Shapes indicate the four different sites and the colours the horizon. Regression lines are illustrated by generalized linear smoothing and the shaded areas indicate the 95% confidence interval.

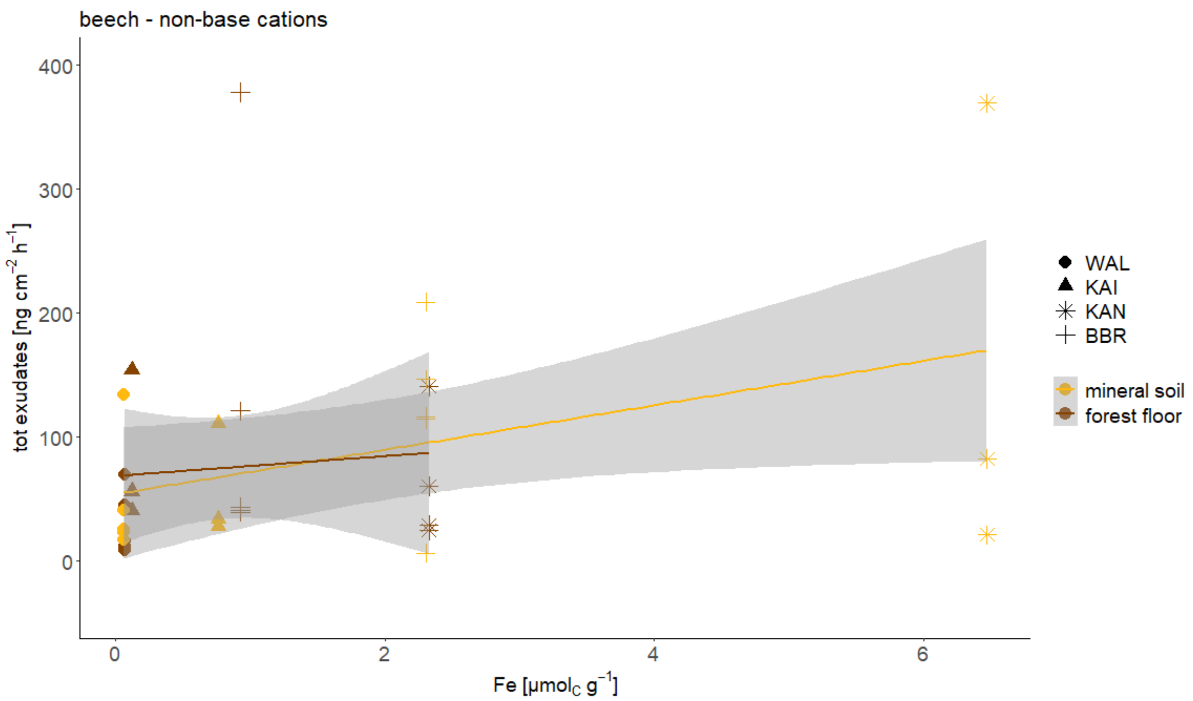


***Figure S6***: Total exudates in relation to exchangeable Fe concentration for beech. Shapes indicate the four different sites and the colours the horizon. Regression lines are illustrated by generalized linear smoothing and the shaded areas indicate the 95% confidence interval.

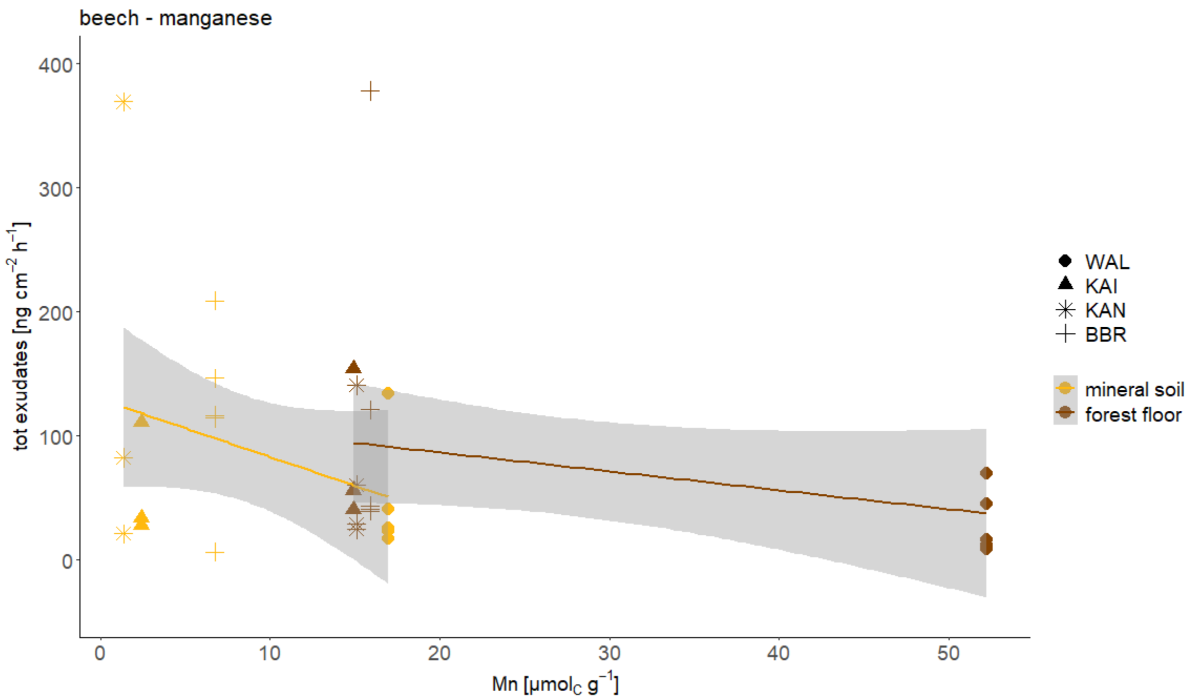


***Figure S7***: Total exudates in relation to exchangeable Mn concentration for beech. Shapes indicate the four different sites and the colours the horizon. Regression lines are illustrated by generalized linear smoothing and the shaded areas indicate the 95% confidence interval.

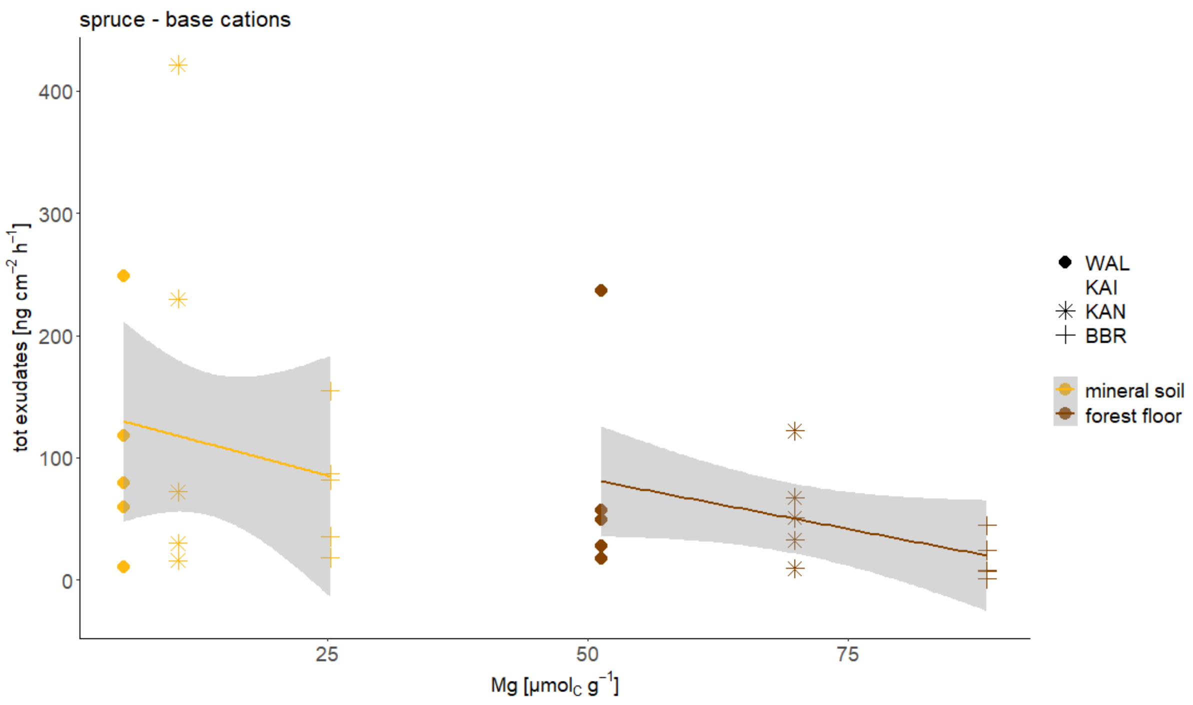


***Figure S8***: Total exudates in relation to exchangeable Mg concentration for spruce. Shapes indicate the four different sites and the colours the horizon. Regression lines are illustrated by generalized linear smoothing and the shaded areas indicate the 95% confidence interval.

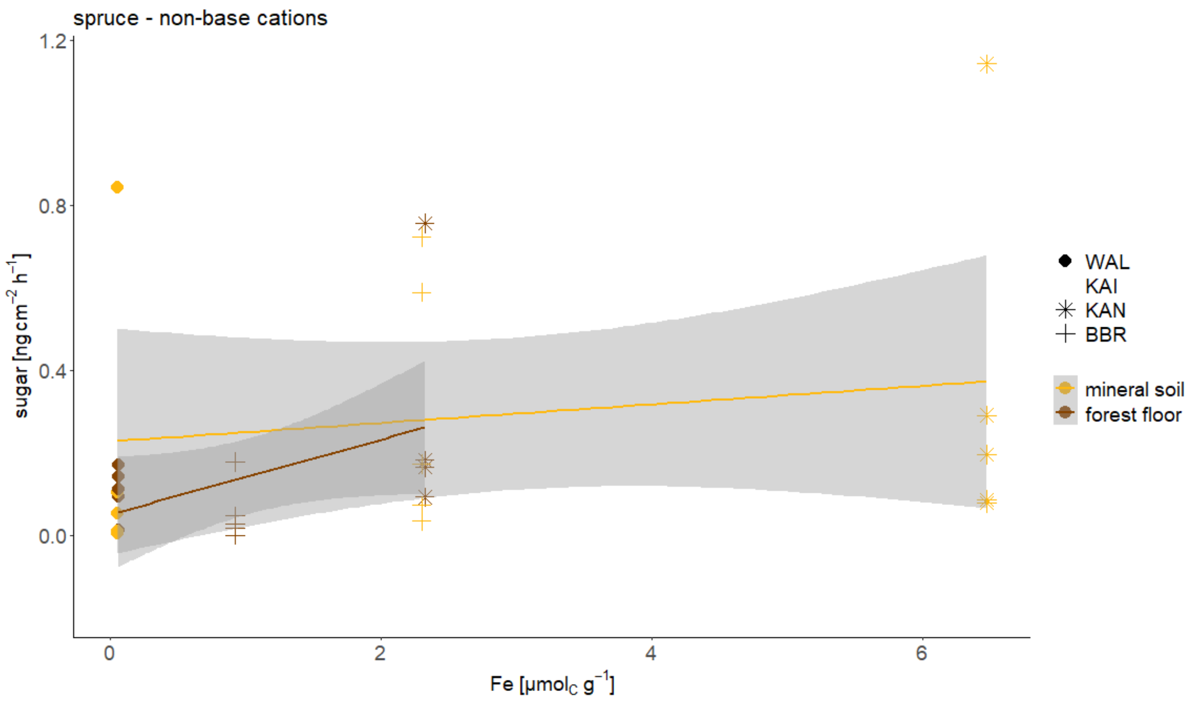


***Figure S9***: Sugar exudation in relation to exchangeable Fe concentration for spruce. Shapes indicate the four different sites and the colours the horizon. Regression lines are illustrated by generalized linear smoothing and the shaded areas indicate the 95% confidence interval.

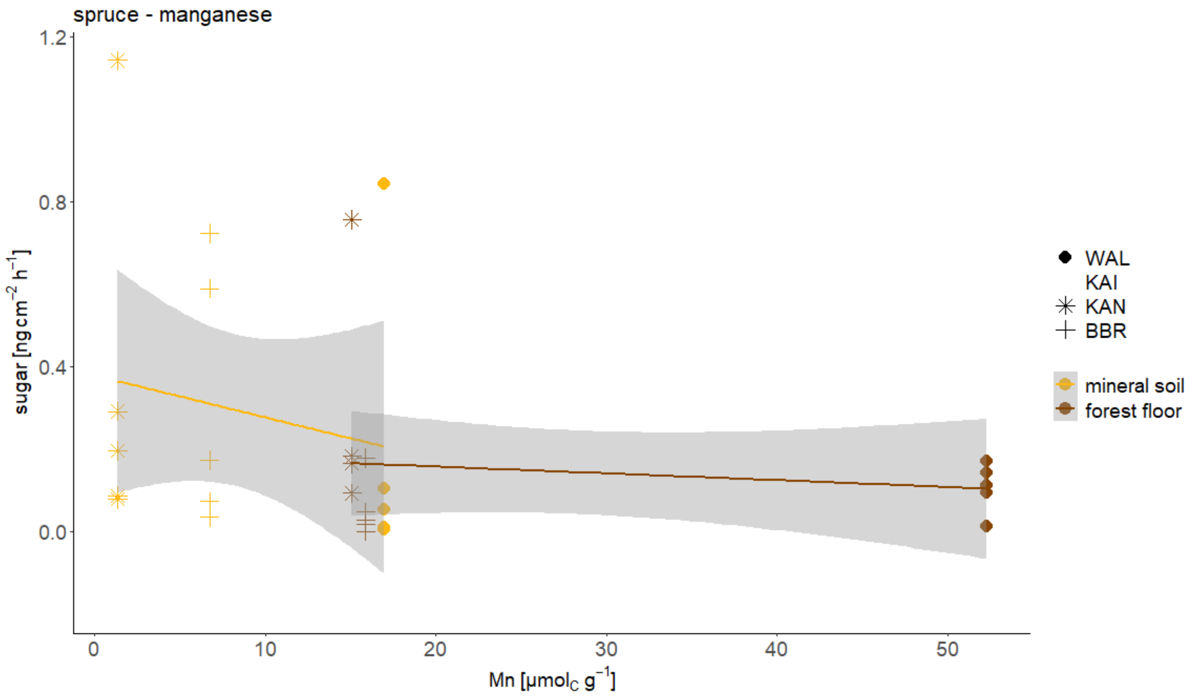


***Figure S10***: Sugar exudation in relation to exchangeable Mn concentration for spruce. Shapes indicate the four different sites and the colours the horizon. Regression lines are illustrated by generalized linear smoothing and the shaded areas indicate the 95% confidence interval.
